# Allosteric communication between ligand binding domains modulates substrate inhibition in adenylate kinase

**DOI:** 10.1101/2022.11.21.517316

**Authors:** David Scheerer, Bharat V. Adkar, Sanchari Bhattacharyya, Dorit Levy, Marija Iljina, Inbal Riven, Orly Dym, Gilad Haran, Eugene I. Shakhnovich

## Abstract

Enzymes play a vital role in life processes; they control chemical reactions and allow functional cycles to be synchronized. Many enzymes harness large-scale motions of their domains to achieve tremendous catalytic prowess and high selectivity for specific substrates. One outstanding example is provided by the three-domain enzyme adenylate kinase (AK), which catalyzes phosphotransfer between ATP to AMP. Here we study the phenomenon of substrate inhibition by AMP and its correlation with domain motions. Using single-molecule FRET spectroscopy, we show that AMP does not block access to the ATP binding site, neither by competitive binding to the ATP cognate site nor by directly closing the LID domain. Instead, inhibitory concentrations of AMP lead to a faster and more cooperative domain closure by ATP, leading in turn to an increased population of the closed state. The effect of AMP binding can be modulated through mutations throughout the structure of the enzyme, as shown by the screening of an extensive AK mutant library. Mutation of multiple conserved residues leads to increased substrate inhibition, suggesting a positive selection during evolution. Combining these insights, we developed a model that explains the complex activity of AK, particularly substrate inhibition, based on the experimentally observed opening and closing rates. Notably, the model indicates that the catalytic power is affected by the microsecond balance between the open and closed states of the enzyme. Our findings highlight the crucial role of protein motions in enzymatic activity.

**Significance Statement:** How conformational dynamics affect the catalytic activity of enzymes remains a topic of active debate. We focus here on the domain closure dynamics of adenylate kinase (AK) and how they are affected by substrate inhibition. By screening an extensive mutant library, we show that this feature of the enzyme is well conserved in evolution. Importantly, domain closure is required in order to bring AK’s substrates close together for their chemical reaction; single-molecule FRET studies directly measure the populations of the open and closed states. We find that overpopulation of the closed state can be detrimental to activity. The results allow us to develop a kinetic model that properly accounts for AK kinetics by combining conformational dynamics and biochemical steps.

## INTRODUCTION

Enzymes have been designed by nature to accelerate vital chemical reactions by many orders of magnitude. Quite a few of them have evolved to harness large-scale motions of domains and subunits to promote their activity. The study of the structural dynamics of these proteins is essential to decipher how they function and are regulated, as has been demonstrated in multiple experimental and theoretical reports (1–4). The enzyme adenylate kinase (AK) is a paradigmatic example of a strong relation between conformational dynamics and catalytic activity (5–8). AK plays a key role in maintaining ATP levels in cells by catalyzing the reaction ATP + AMP ⇄ ADP + ADP (9, 10). It consists of three domains: the large CORE domain, the LID domain that binds ATP, and the nucleotide monophosphate (NMP)-binding domain that binds AMP. X-ray crystallographic studies showed that the LID and NMP domains undergo a major conformational rearrangement upon substrate binding (Figure 1a) (11–13). This movement, termed domain closure, forms the enzyme’s active center and helps to exclude solvent molecules that might interfere with the chemical reaction. AK’s domain-closure dynamics have been studied using NMR spectroscopy (6–8, 14) and fluorescence experiments (5, 6, 15), as well as multiple molecular dynamics simulations (16–21). In particular, our recent single-molecule FRET (smFRET) experiments demonstrated that AK’s opening and closing rates are significantly faster than previously reported (22). In the presence of substrates, domain closure is completed in just a few tens of microseconds. These are likely the fastest large-scale conformational dynamics measured to date, and they are orders of magnitude faster than the overall turnover rate of the enzyme. It was proposed that these fast domain movements might assist the enzyme in orienting its substrates optimally for catalysis, a hypothesis supported by recent simulations (23).

**Figure 1.**
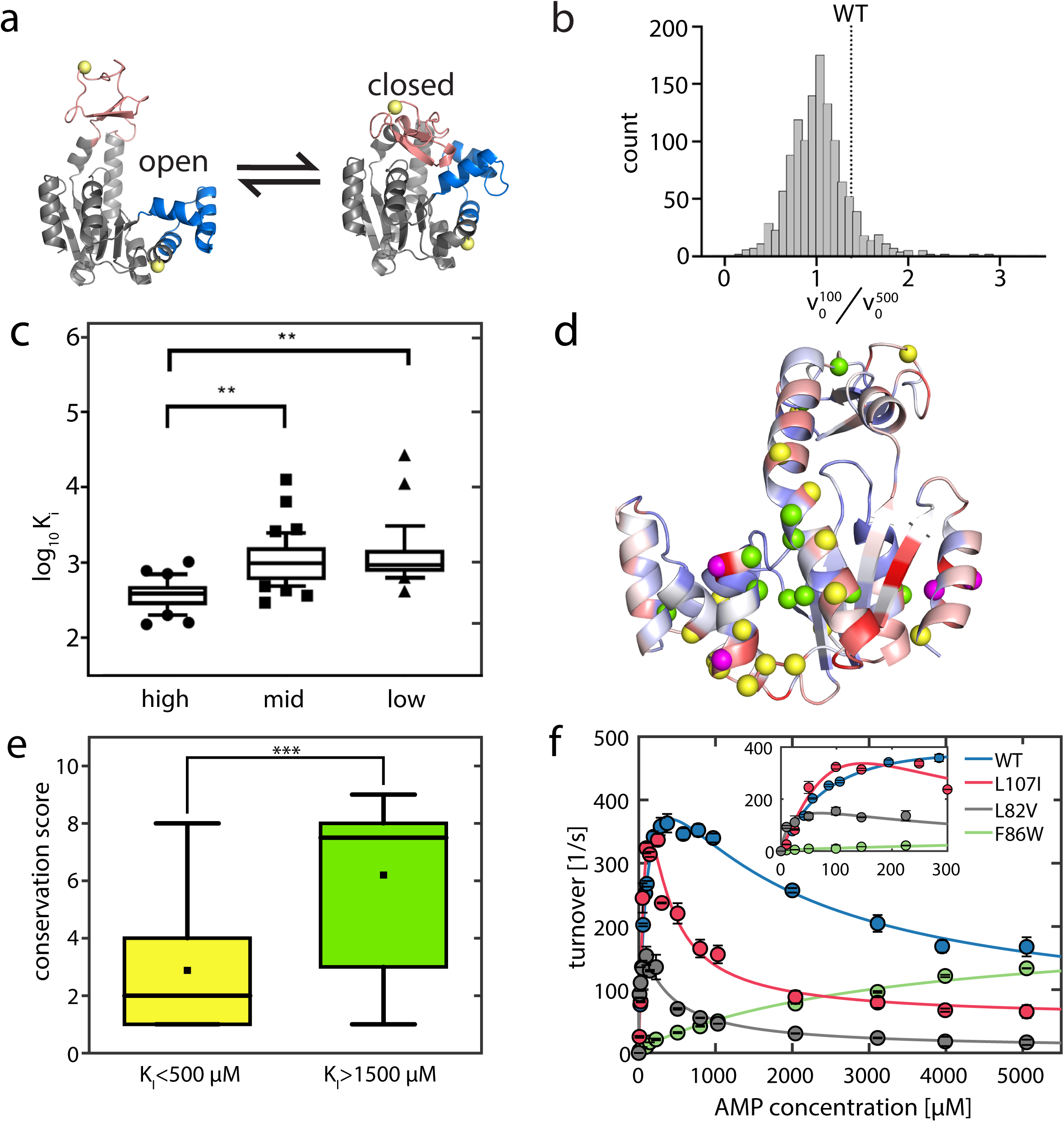
Structural and kinetic attributes of AK variants. (a) Structure of AK with its three domains. The CORE domain (grey) connects the LID domain (pink) and the NMP domain (blue). The yellow spheres indicate the positions for the attachment of fluorescent dyes. The protein can undergo a conformational change from the open state (left, PDB 4AKE) towards the closed conformation (right, PDB 1AKE). b) Distribution of in-lysate activity of 1248 colonies in terms of the screening parameter 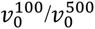. WT corresponds to a 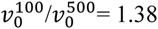. c) Distribution of *K_I_* values of purified proteins from library variants with high (top 64), mid (64 around median) and low (bottom 64) values of 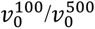 measured in-lysate. (d) Conservation score mapped onto the open state with high to low conservation represented as blue to red. The positions with mutants having a *K_I_* <=500 μM for AMP are shown as yellow spheres, while those with *K_I_* >=1500 μM are shown in green. The four magenta spheres are positions where both inhibited and uninhibited mutations are observed (see Table S1). The conservation scores were obtained from the ConSurf web server (28, 29). (e) The distributions of the conservation scores in the strong and weak inhibition bins were found to be statistically different (p<0.01, Student’s t-test). The mutations that result in strong inhibition (*K_I_* <=500) usually occur at less conserved positions (2.9±0.6), while inhibition-relieving mutations (*K_I_* >=1500) occur mostly at highly conserved positions (6.2±0.9). (f) Activity curves of WT AK and selected mutants show increased SI (L107I, L82V) or loss of inhibition (F86W).

Interestingly, AK also incurs a pronounced AMP-mediated substrate inhibition (SI) (24, 25). Despite many efforts, the mechanism behind SI has not been elucidated convincingly. While some studies have hinted toward the competitive binding of AMP at the ATP site (26), others claimed that AMP substrate inhibition in AK is uncompetitive in relation to ATP (24). Whether large-scale domain motions in AK relate to SI has also been debated. Bulk FRET studies have indicated that inhibition arises due to substantial closing of the LID domain upon AMP binding (25). However, it was unclear if this effect was induced by direct binding of AMP to the ATP site (competitive inhibition at the ATP site) or through allosteric modulation (i.e., AMP binding at its cognate site affects LID domain movement). Other studies suggested that at high AMP concentrations, binding of AMP before the release of ADP from the LID domain might lead to the formation of an abortive complex that causes inhibition (24). The latter effect could arise if LID opening is slow and is the rate-limiting step in catalysis, as has been proposed (6, 14, 15).

In this work, we carried out experimental and computational studies on several inhibition-prone as well as non-inhibited mutants of AK (27) to determine the mechanism of AMP-mediated SI in AK. Based on the screening of an extensive AK mutant library, we show that mutations throughout the enzyme’s structure modulate SI, suggesting that it has been under positive selection during evolution. We establish that inhibition cannot be explained by either competitive binding of AMP at the ATP site or spurious LID domain closure by AMP in the absence of ATP. Instead, we find that for inhibition-prone variants of AK, AMP facilitates domain closure at lower concentrations of ATP, which may interfere with proper substrate binding mechanics required for the reaction. Based on these results, we propose a kinetic model that combines conformational dynamics and chemical steps to explain the enzymatic activity of AK with substrate inhibition.

## RESULTS

### Structural and kinetic attributes of AK variants with substrate inhibition

In an earlier study (27), we characterized a series of mutants of AK and found a correlation between AMP-mediated SI and protein stability. An important question that arises is whether there is a specific region that is responsible for the gain or loss of SI, possibly by affecting how the substrates bind to the protein. To answer this, we created an extensive library of AK mutants and characterized their SI profiles. We particularly sought to identify mutations that alter SI without perturbing the enzymatic activity. Accordingly, using the crystal structure of AK with the bound inhibitor AP_5_A (12), we first identified 71 residues that were at least 8Å away from the inhibitor and their sidechains were not involved in any interactions with the rest of the protein (Figure S1a, see details in Methods). We then generated a library of 923 mutants wherein 13 different amino acids were introduced at each of these 71 locations identified above (see Methods). Next, we developed a screening method (Figure S1b) to measure the extent of AMP-mediated SI directly from the cell lysate of 1248 individual colonies by measuring the ratio of the initial reaction velocity at two different concentrations of AMP (100 µM and 500 μM, Figure 1b).

We selected ~200 candidates from this dataset that showed lower, higher, and similar SI as the WT protein and sequenced them by Sanger sequencing to recover 86 unique mutations with clean chromatograms in the first pass. We further purified these proteins and measured their activity profiles (Figure 1c). For a quantitative comparison between the mutants and literature values, the AMP-related Michaelis constant (*K_M_*) and inhibition constant (*K_I_*) were obtained by a fit to a model of uncompetitive inhibition (see Methods), and the values are given in Table S1. We mapped the positions of the inhibition-prone mutants (*K_I_* =< 500) and the non-inhibited mutants (*K_I_* >= 1500) on the structure of AK (Figure 1d). Both effects can be observed for residues distributed all over the structure rather than in specific regions of the protein. As shown in Figure 1e, we observed that at those locations where mutation led to a loss of inhibition, the WT residue had a statistically significant (p<0.01, Student’s t-test) higher conservation than those that led to an increase of inhibition. (The conservation score was determined by a high frequency of occurrence in a multiple sequence alignment (28, 29)). In other words, the gain of SI upon mutation tends to occur at positions of low conservation, while the loss of SI occurs at highly conserved locations. This signifies that SI in AK has been under positive selection during evolution. Since back-to-consensus mutations are generally stabilizing (30), this plot also signifies that higher protein stability generally leads to more SI in AK, as seen from our previous study (27).

The mutant screening experiment did not identify a specific structural attribute that is responsible for SI by AMP. Further, a crystallographic study of one of the mutants also did not demonstrate a structural change that may explain the effect of AMP. In particular, we determined the crystal structure of L107I AK, which shows strong inhibition, in complex with Ap_5_A to a resolution of 2.05 Å. The structure was found to be very similar to other closed AK structures (12, 31, 32) (Figure S2), with a 0.3 Å root-mean-square deviation (RMSD) from the 1AKE structure (12). As no differences in the static structure of AK were found that could explain differences in enzymatic activity, these are more likely rooted in dynamic interactions. Therefore, we turned to studies of the dynamic properties of AK mutants in relation to SI.

### From biochemistry to structural dynamics

To explain the origins of SI in AK and similar multi-substrate enzymes, previous studies have primarily considered three hypotheses. The first involved binding of the inhibiting substrate to an additional site (e.g. the ATP site) (33). The second, uncompetitive inhibition by AMP through blocking the accessibility of the ATP binding site (24, 25). Finally, it was suggested that kinetic differences between alternating pathways towards the fully substrate-bound complex could lead to inhibition (34).

To probe the origins of SI further, we selected three variants of AK with similar thermal stabilities as the WT: L82V and L107I (27), which are more inhibited than the WT, and F86W, which is not inhibited by AMP (24). Michaelis-Menten curves for these variants as a function of AMP concentration are shown in Figure 1f. We first investigated if SI in AK arises due to AMP having a second binding site in addition to the one on the NMP domain. The additional binding site could be the ATP-binding site – as AMP is a close structural analog of ATP – or any other yet-unknown site. In that context, we found that for the WT (Figure S3a, Table S2), the presence of excess ATP led to a gradual and dose-dependent loss in SI, which is a classic apparent sign of competitive inhibition at the ATP binding site. In other words, it seemed possible that the competitive binding of AMP at the ATP binding site at high concentrations causes inhibition, which is relieved by excess ATP in the solution. If this were true, then under conditions when the AMP concentration vastly surpasses the ATP concentration, binding of AMP to the ATP site would be preferred and result in a complete loss of activity. Contrary to this expectation, the enzyme retains significant residual activity even at high AMP concentrations (5, 24, 25), as seen particularly well for L107I (Figures 1f, S3b).

We therefore ask: does AMP inhibit AK by binding to the ATP binding site in the LID domain, or does it affect the catalytic properties differently? To gain insight into this question, we turned to smFRET spectroscopy to measure the domain movements directly. AK was labeled at residues 73 and 142, positioned in the CORE and LID domains, respectively (Figure 1a), and studied in solution (22). In the absence of substrates, AK molecules yielded a FRET efficiency histogram dominated by a peak at ~0.4 (Figure 2a, blue), corresponding to the open conformation. A shoulder towards higher FRET efficiency values indicated that the closed conformation is also sampled, though to a small extent (6, 22, 35). The addition of ATP (Figure 2a, green and purple) shifted the histogram towards higher FRET efficiency values, resulting from a considerable increase in the population of the closed conformation. In the presence of 1 mM AMP (orange), similar shifts were observed, but they required much lower concentrations of ATP.

**Figure 2.**
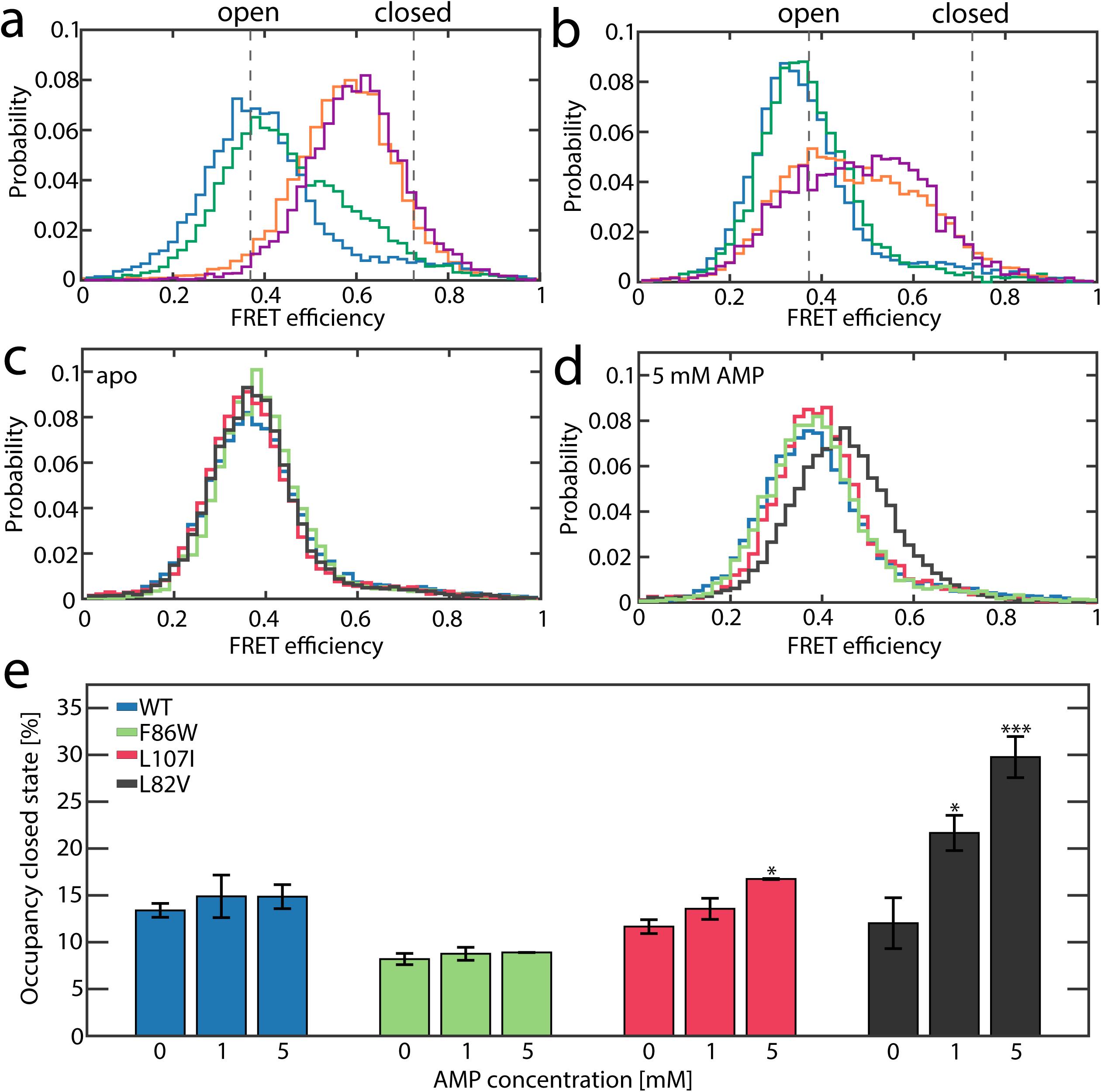
AMP does not close the LID domain. a-b) FRET efficiency histograms of the WT protein. (a) The apoprotein (blue trace) mainly adopts an open conformation, occasionally exploring the closed state. The binding of ATP increases the population of the closed state, as demonstrated by histograms at 100 μM (green) and 1 mM (purple) ATP. When 1 mM AMP is added, a significant population of the closed state is attained even at a low ATP concentration of 50 µM (orange). In this experiment, ADP was also added at concentrations that maintained the equilibrium of the enzymatic reaction (see Methods). The dashed lines indicate the most likely positions of the open and closed state (0.37/0.72 for the WT, respectively). (b) AMP alone, even at 30 mM (green), leads to very minimal changes in FRET efficiency compared to the apoprotein (blue). ATP-γ-S, a slowly hydrolyzable analog of ATP, binds to the ATP binding site and triggers partial closure at 50 µM (orange). In the presence of both ligands (purple), ATP-γ-S is still bound to the ATP-binding site and not outcompeted by AMP, which would trigger LID opening. Instead, the binding of AMP to its own site even promotes closure to a minor extent. (c) For the apoprotein, the FRET efficiency histogram of the WT (blue) and all mutants are similar. Shown are F86W (green), L107 (red) and L82V (black). (d) FRET efficiency histogram for all mutants in the presence of 5 mM AMP. Only for L82V, a significant shift to higher FRET efficiency is detected. e) Values for the occupancy of the closed state as a function of AMP concentration were obtained by H^2^MM analysis. The experiment confirms that the occupancy of the closed state for the WT (blue), L107I (red) and F86W (green) is not significantly altered by AMP alone. In contrast, for L82V (black), a partial closure is observed. Asterisks indicate the significance of the deviation of parameters from the apoprotein (***: p<0.01, **: p<0.05, *: p<0.1, no index: p>0.1, Student’s t-test). Error bars are given as the standard error of the mean based on 3 independently prepared samples.

Surprisingly, in the presence of high concentrations of AMP as a sole substrate, the FRET efficiency histogram of the WT protein remained unchanged (Figure 2b), indicating that AMP alone cannot cause LID domain closure. To see if the effect of AMP is different in mutants with stronger or weaker SI, we also double-labeled the three variants introduced above (L107I, L82V and F86W). None of these single-point mutants significantly affected the FRET efficiency histogram of the apoenzyme (Figure 2c), and the addition of AMP alone did not induce a significant shift (Figure 2d). The only exception was L82V, the most inhibited mutant, which showed a slight shift towards higher FRET efficiencies. To accurately retrieve changes in the populations of the open and closed conformations of the enzyme under various conditions, we used the hidden-Markov model-based analysis method H^2^MM (36). This algorithm employs photon arrival times as input for an optimization process that obtains both occupancies and rates of exchange of conformational states and is capable of retrieving microsecond dynamics. Representative trajectories including state assignments using the Viterbi algorithm are shown in Figure S4. The model we used involved only two conformational states, open and closed (Figure 1a). The resulting parameters were validated with different tests as outlined in the Supporting Information (Figure S5-8, Table S3+4). A notable increase in the population of the closed state by AMP was seen only for L82V (Figure 2e). Interestingly, both closing (Figure S9a) and opening (Figure S9b) rates were accelerated in the binary AK-AMP complex. This effect was particularly substantial for L107I and L82V. In contrast, no changes were seen for the uninhibited F86W.

Clearly, AMP alone does not promote the closed state of the LID domain in AK. However, this does not rule out that AMP binds to the ATP site and blocks access for the native substrate, thereby inhibiting the protein. To check this point, we tested whether AMP can competitively replace ATP from its cognate binding site. To limit the potential effect of ADP, which binds to both the ATP and AMP site, we utilized the slowly hydrolyzable ATP analog ATP-γ-S. In the presence of 50 µM ATP-γ-S, partial closure was observed (Figure 2b). If AMP can displace ATP/ATP-γ-S from its native binding site, adding a vast excess of AMP should induce a shift towards the open conformation. Instead, we observed a minor shift towards the closed state in the FRET efficiency histograms, rendering the possibility that AMP acts as a competitive substrate inhibitor very unlikely.

To provide support to the point that AMP does not bind at the ATP site, in addition to the single-molecule studies, we rationally designed a mutant Q92Y that is predicted to weaken AMP binding at its cognate site, based on the structure of *E. coli* AK with bound Ap_5_A (12). The mutation indeed led to a loss in activity for both the WT (Figure S10a) and L82V (Figure S10b). Using Differential Scanning Fluorimetry to obtain the change in thermal melting temperatures of AK upon ligand binding as a function of ligand concentration, we found that Q92Y on the background of WT weakens AMP binding (Figure S10c). On the background of the highly inhibited mutant L82V (L82V-Q92Y mutant), it completely abolished AMP binding (Figure S10d), while ATP binding was retained for both WT and L82V backgrounds (Figure S10e-f). This experiment indicated that AMP cannot bind elsewhere to its cognate site. This was also confirmed by isothermal titration calorimetry studies, which found no more than one binding site for both ATP and AMP (37).

### AMP binding has an allosteric effect on the LID domain

Given the observed effect of AMP binding at its own site on LID domain dynamics, it is imperative to find whether AMP further modulates the effect of ATP on domain closure. Figure 3 shows how the population of the closed state of the enzyme changes in response to substrate binding to the LID domain, either with ATP alone or also in the presence of 1 mM AMP. (In the latter case, ADP was also added to maintain the enzymatic reaction under equilibrium-see Methods and Table S5). The very presence of ATP as a sole substrate (Figure 3, circles) enabled a large conformational change from a predominantly open to a more closed conformation. The maximum occupancy that can be achieved for the closed state is 56-78%, in agreement with previous studies (22, 38). Fitting the curves in Figure 3 to simple binding isotherms provided the transition midpoint concentrations (which we call *C_50_)*, which were found to correlate with the strength of substrate inhibition (Table S6). The impact of a fixed concentration of AMP on the ATP-dependent population of the closed state is shown in squares in Figure 3. LID domain closure occurred at a 20-40 times lower substrate concentration in the presence of AMP (Table S6), indicating remarkable cooperativity in substrate binding. This cooperativity was almost entirely absent for the non-inhibited mutant F86W (Figure 3d). Interestingly, previous work has shown that AMP binding reduces the *K_M_* for ATP in the WT but not in the F86W mutant (24). The strongest cooperativity was observed for the most inhibited mutant, L82V (Figure 3c). Furthermore, in this mutant, the closed state was significantly more populated (78.4±0.7%) than for the WT (57.1±0.6%).

**Figure 3.**
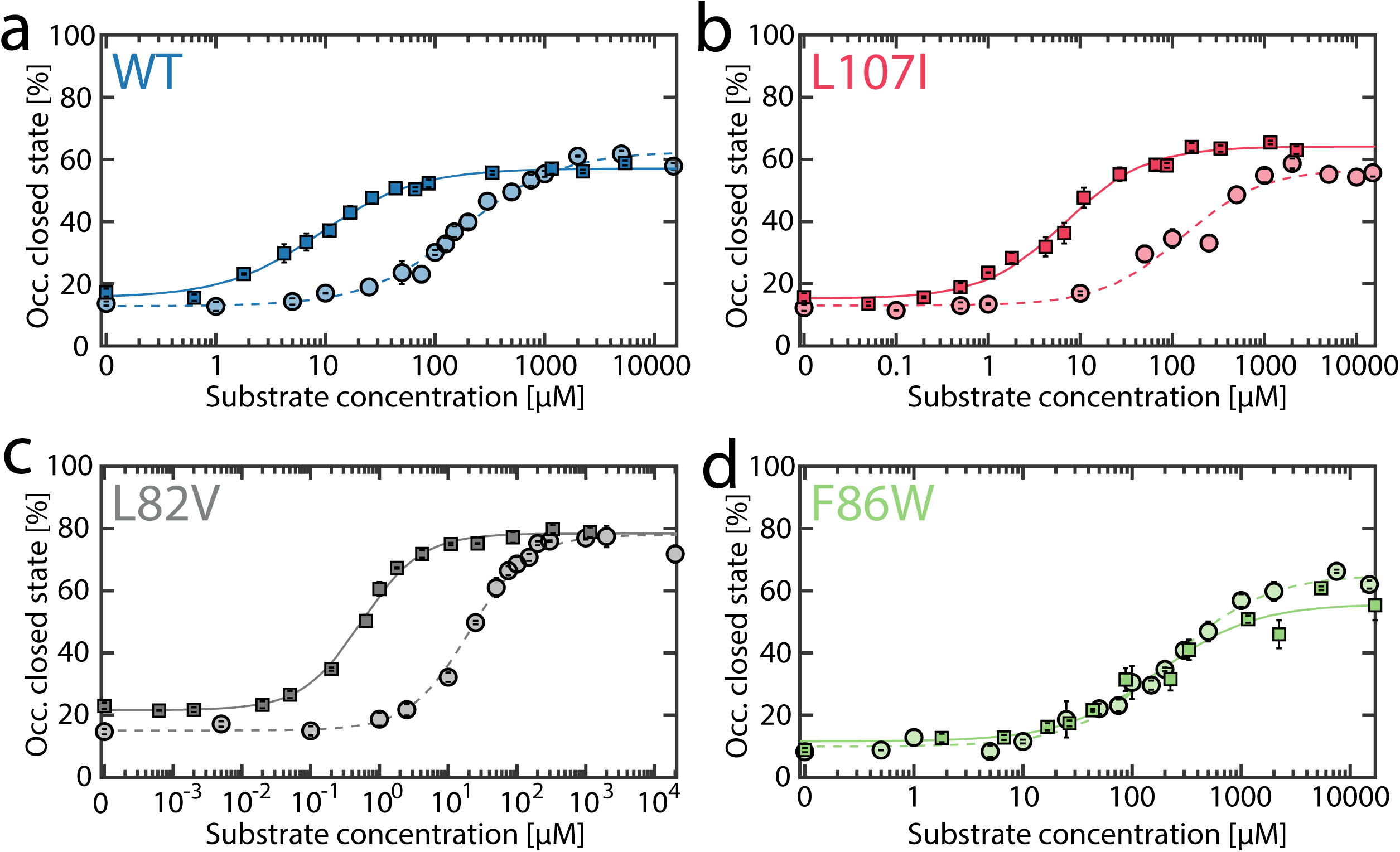
ATP-dependent closure of the LID domain. The occupancy of the closed state was determined by H^2^MM in the presence of either solely ATP (circles) or also 1 mM AMP and appropriate concentrations of ADP to maintain equilibrium, as described in Methods (squares). In all variants, the LID domain closes at high ATP levels. For (a) WT, (b) L107I and (c) L82V, the presence of 1 mM AMP promotes closure at much lower ATP concentrations. In the non-inhibited mutant (d) F86W, AMP has no significant effect. Dashed (ATP only) and straight (all substrates) lines indicate fits to a binding model as described in Supplementary Note 1. Error bars are given as the standard error of the mean based on 2-3 independently prepared samples. As in Figure 2a, ADP was added to maintain equilibrium; the substrate concentration on the abscissa is the total concentration of ligands for the ATP binding site (i.e. ATP+ADP, Table S5).

It is clear from the results of Figure 3 that AMP has an effect on the closure of the LID domain, even though it does not directly bind to that domain. AMP must therefore affect the LID domain allosterically. To shed further light on the effect of AMP on LID domain mechanics, we studied domain-closure dynamics at a fixed concentration of ATP (1 mM, together with specified concentrations of ADP to maintain equilibrium) and increased concentrations of AMP (Figure 4). As reported earlier (22), both opening and closing rates are much faster than the enzymatic turnover (Table S7). In the WT protein in the absence of AMP (Figure 4a), closing rates (27,307±752 s^−1^) were slightly faster than opening rates (23,937±1,123 s^−1^). Strikingly, the gradual addition of AMP did not affect the rate of domain opening. For the closing rate, small AMP concentrations – at which no inhibition is detected (up to ~1 mM) – also did not have a measurable impact. However, at inhibitory AMP concentrations (> 1 mM), domain closure was significantly accelerated (up to 50,289±2,795 s^−1^), leading to a higher occupancy of the closed state of 63.5±0.4% (Figure 4d). Interestingly, this overpopulation of the closed state coincided with the range of AMP concentrations where the most substantial drop in activity (Figure 1f) was observed. A very similar picture was seen with the more inhibited variants (Figure 4b-c). No measurable impact of AMP binding was detected for the opening rates, whereas the closing rates increased above 100 µM. A *C_50_* value of 2568±820 µM was found for the WT, with significantly lower values (~500 µM) for L107I and L82V. Moreover, L107I was shifted further towards the closed state (64.8±0.5%) than it is possible in the presence of ATP alone (55.8±0.8%). For L82V, the population of the closed state (75.5±0.8%) was the largest, which might match that mutant’s overall low activity. No enhancement of the closing rate was detected for the non-inhibited mutant F86W (similar to Figure S9).

**Figure 4:**
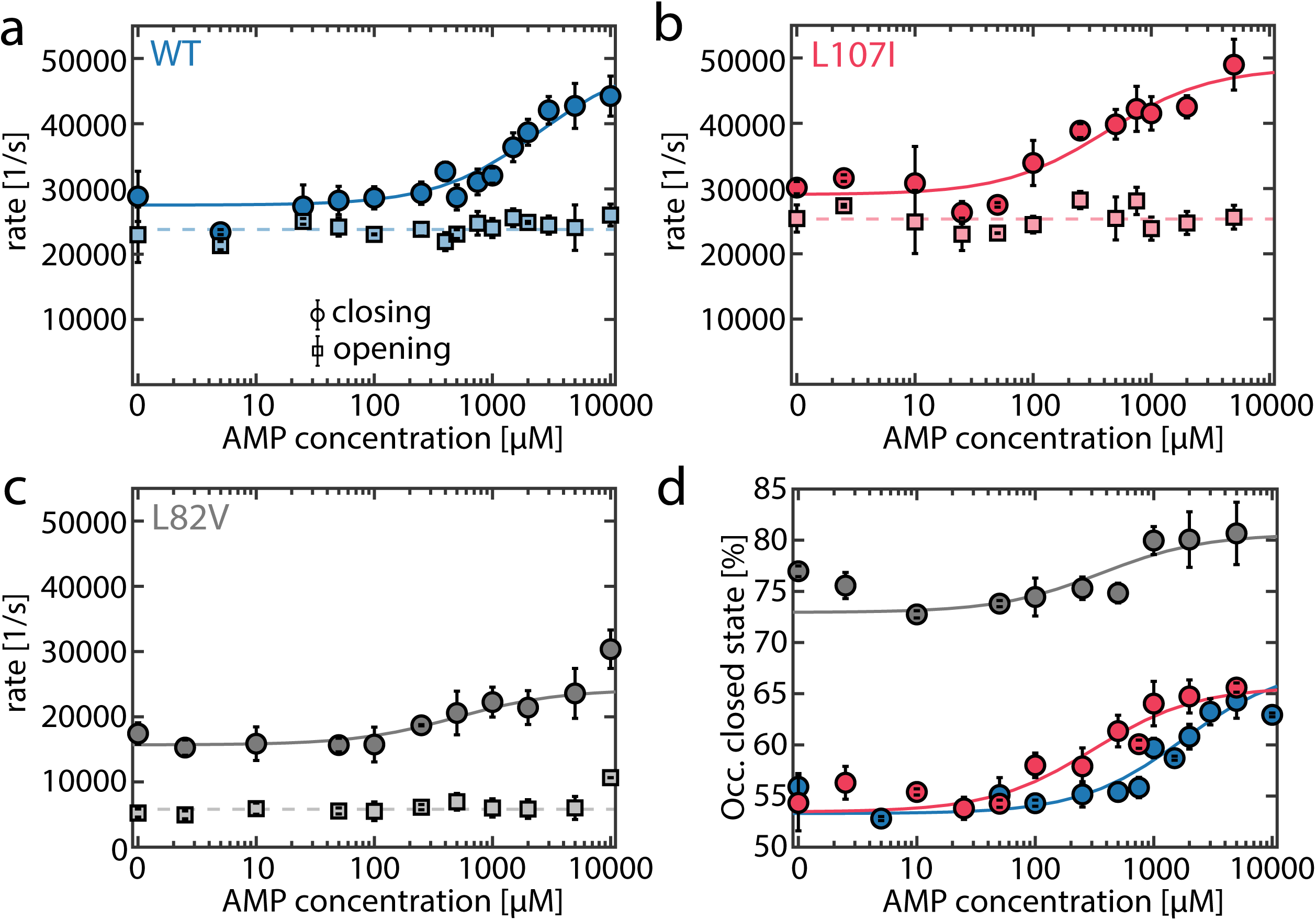
AMP accelerates LID domain closing. a-c) Opening (circles, solid lines) and closing (squares, dashed lines) rates for a) the WT (blue), b) L107I (red) and c) L82V (grey) in the presence of increasing concentrations of AMP and a fixed concentration of ATP (1 mM). Opening rates are not affected by the presence of AMP. The closing rates were fitted to a model described in Supplementary Note 1. d) Occupancy of the closed state based on the rates shown in a-c). Error bars are given as the standard error of the mean based on 2-3 independently prepared samples.

### A model to explain substrate inhibition

Given that AMP inhibits AK’s enzymatic activity and also affects domain-closure dynamics, we postulated that the two phenomena are correlated. Therefore, we devised a kinetic model for AK, based on the hypothesis that different closed states can be sampled, depending upon the order of ligand binding. Some of these states are incompatible with catalysis and must convert to the catalysis-prone state before the reaction occurs. As per our model (Figure 5a), the apoenzyme E can be present in both open (E^O^, blue) and closed (E^C^, red) states. (States are defined only for the ATP-binding LID domain.) The status of the LID domain does not affect the binding of AMP to the NMP domain; hence, both open and closed states can bind AMP. However, ATP can bind only to the open state.

**Figure 5:**
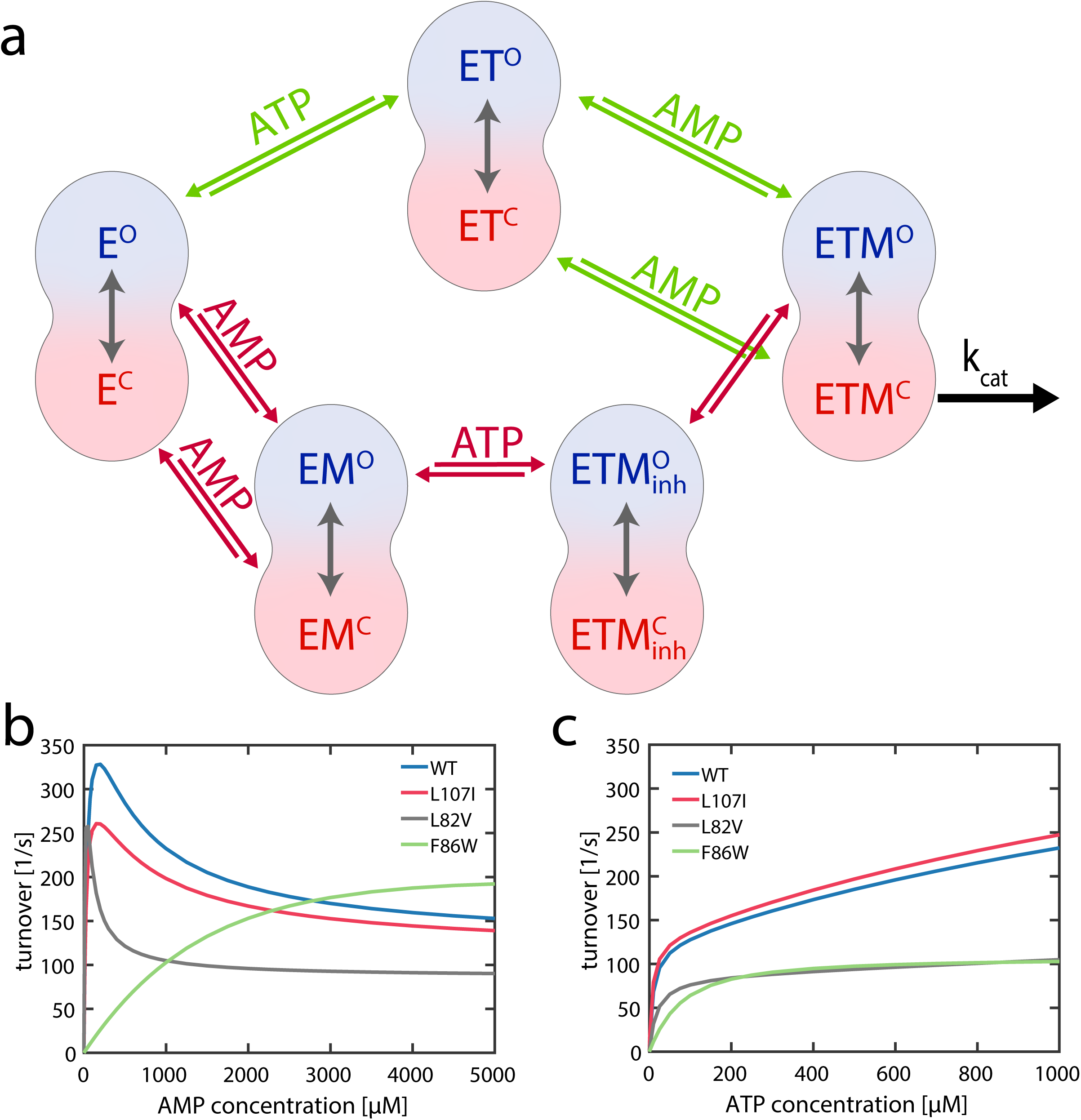
A model to explain substrate inhibition. (a) Scheme of the model, which is based on two competing pathways: When ATP binds first (green pathway) to the apoenzyme (E), this results in the ATP-bound state ET. Subsequent binding of AMP leads to a productive closed state (ETM) that undergoes the chemical step of the enzyme in the closed conformation (ETM^C^). However, if AMP binds first (red pathway, via EM), subsequent binding of ATP causes the formation of ETM_inh_, which deviates from the productive state ETM in the orientation of the substrates. It is assumed that the closed state 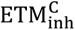 does not enable phosphotransfer due to an inappropriate substrate orientation. The corresponding open conformation 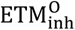 must convert to ETM^O^ to enable catalysis. Binding or reorientation of ATP, which lead to a productive catalytic conformation, cannot occur in the closed conformation (23). (b-c) Simulated activity curves of WT and mutants as a function of (b) AMP and c) ATP concentrations based on the model (a) and experimentally observed parameters of opening and closing rates. The rate of conversion from ETM1^O^ to ETM2^O^ was fixed at 250 s^−1^. The model shows AMP-mediated SI for WT, L107I and L82V, but not for F86W. No SI was observed as a function of ATP concentration, in accordance with experimental data.

If ATP binds first at the LID domain, the ET^O^ and ET^C^, i.e. open and closed conformations of the ATP-bound protein, respectively, are formed. Both of these states can subsequently bind AMP and form a double substrate-bound state, ETM. If AMP binds first at the NMP domain to form EM^O^ and EM^C^, the consequent binding of ATP leads to the formation of ETM_inh_. ETM and ETM_inh_ are distinguished by their binding order – ATP first, then AMP or vice versa-based on a hypothesis that initial AMP binding may result in a limited or different sampling of the relative orientation of the nucleotides (see below). Hence, 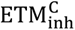 is an unproductive closed state, while ETM^C^ is productive. The two ETM states can interconvert only through their open states, as suggested recently by MD simulations (23). At the same time, phosphoryl transfer can happen only through ETM^C^. In essence, we have introduced a kinetic hindrance for the path that binds AMP first. When a typical activity assay of AK is carried out in the bulk in the presence of saturating ATP concentrations, at low AMP levels greater flux of the reaction goes through the pathway involving the productive ETM intermediate. However, as AMP levels increase, a larger fraction of the flux goes through the pathway involving the unproductive ETM_inh_ state, leading to a drop in activity.

To simulate kinetics based on this model, we used experimentally observed opening and closing rates of the LID domain for E (E^O^ ⇌ E^C^), ET (ET^O^ ⇌ ET^C^), EM (EM^O^ ⇌ EM^C^), ETM (ETM^O^ ⇌ ETM^C^) as input in the model (Table S8). We assumed that each species has one specific opening and closing rate. The concentration of nucleotides determined the occupancy of each of these states (22). The single-molecule experiments could not distinguish between ET and ETM, as no change in the apparent protein dynamics was observed upon the addition of AMP at non-inhibiting concentrations (Figure 4). Therefore, the rates adopted for these two states were identical. The rates for the ETM_inh_ state (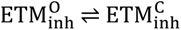) were derived from asymptotic values obtained from the binding isotherm fits of Figure 4 (Supplementary Note 1). We also assumed a single rate constant for the actual phosphotransfer step (500 s^−1^) and the conversion rate between 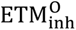 and ETM^O^ (250 s^−1^) for all mutants. Nucleotide binding rates are given in Table S9.

Under these model assumptions, a simulation of the enzyme activity at a saturating ATP concentration (1mM) and varying AMP concentrations (Figure 5b) was effectively able to qualitatively reproduce the SI pattern of the WT as well as the mutants L107I and L82V. Further, no inhibition was observed for the F86W mutant, as expected. On the other hand, using a saturating AMP concentration (1mM) and varying ATP concentrations, the simulation did not show SI for any mutant (Figure 5c). The ability to obtain the correct level of inhibition by AMP for the different mutants provides a strong support for the model. While in principle we could use the model to quantitatively fit the enzyme activity curves, we believe that our lack of knowledge on some of the involved parameters does not currently warrant such a calculation. Future studies might shed light on several of the unknown parameters, facilitating a more quantitative analysis.

We used the model to obtain some further insight into the role of domain closure dynamics in enzyme activity. To this end, we altered the rates of LID domain closing of each of the ETM states in turn, using one of two ways (Figure S11): either by preserving the open/closed ratio *K*_C_ or by preserving the opening rate, similar to the experimentally observed effect of AMP. In both scenarios, all other parameters were kept constant. As long as the equilibrium ratio *K*_C_ was preserved (Figure S11a), varying the closing rate towards the unproductive 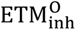 did not affect activity. With *K*_C_ preserved, also varying the rate towards the productive ETM^C^ had a minor effect (Figure S11b). A much more substantial impact of varying the closing rate towards 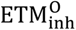 was observed when the opening rate was fixed (Figure S11c). When the conformational equilibrium was altered, the unproductive states became more populated, which caused stronger inhibition. In contrast, for the productive closing (Figure S11d), enhancing the domain closure rate had a positive effect on turnover.

An important criterion determining SI strength is the conversion rate between 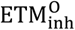 and ETM^O^. Figure S11e depicts how an increase in this rate can almost completely thwart SI while a reduction in the rate increases the effect. In our simulations (Figure 5b/c), we assumed a constant interconversion rate for all mutants, 250 s^−1^. Still, the potent inhibition observed experimentally for L107I and L82V might also be influenced by a slower transition between 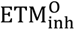 and ETM^O^ for these mutants, and also by other parameters whose accurate values cannot be determined currently.

## DISCUSSION

Despite its comparatively small size, AK’s ubiquity and central role in cellular metabolism have attracted numerous studies. In particular, the complex interplay between conformational changes and enzymatic activity has been frequently investigated. An important aspect here is the AMP-mediated SI, which has been studied for several decades, but no mechanism has been conclusively established. In a previous work, several stabilizing and destabilizing point mutations were introduced into AK, and the mutants exhibited many extremes of SI (from very strong to no SI at all) (27). In the present work, we used several of those mutants (covering a range of SI behavior) to pinpoint a mechanism for inhibition. We show that neither uncompetitive nor competitive binding of AMP can explain SI in AK. Instead, we find that inhibition arises due to strong allosteric communication between the NMP and LID domains, which manifests in a highly cooperative closing of the LID domain for inhibition-prone mutants in the presence of both ligands (ATP and AMP) as compared to ATP alone. Based on this, we propose a model for the enzymatic activity that explains why greater sampling of the LID closed state in the presence of AMP and ATP might lead to SI. According to this model, the order of substrate binding matters. In particular, the initial binding of AMP followed by ATP leads to a closed state that does not allow the correct positioning of the ligands for effective phosphate transfer. Using the experimentally observed opening and closing rates for each state of the enzyme, this model can effectively reproduce the SI profiles of the different mutants studied. The enzymatic cycle of AK has traditionally been understood as a random bi-bi mechanism where the two substrates ATP and AMP can bind in any order. Our data shows that AK activity follows a particular class of random reactions: the substrates can bind in any order, but the outcome of the reaction is different along the two pathways.

Among the two inhibition-prone mutants, L82V displayed somewhat different properties than L107I. Being the most inhibition-prone, L82V showed an increase in closed-state occupancy with AMP alone, indicating that the allosteric communication between the LID and NMP sites is the highest in this mutant. Nevertheless, even for this mutant, opening rates are higher than closing rates in the presence of only AMP; hence, structural inaccessibility of the ATP binding site cannot be the reason for SI. In the presence of both AMP and ATP, the closed state occupancy for L82V is the highest among all mutants (78% vs. 63-65% for others), which is likely the reason for the potent inhibition and overall low activity. In contrast, F86W displayed minimal impact of AMP on LID domain dynamics, and likewise, no inhibition.

Controlling the equilibrium between the open and closed state in AK provides a way to regulate enzymatic activity. Modulating this conformational equilibrium by other means, like mutations or osmolytes, was shown to affect enzymatic catalysis directly (22, 39). AMP also appears to be a major regulator of AK activity (39). In this context, we found that urea can lift AMP-mediated inhibition by repopulating the open state. We will focus on this effect in a future study.

The maximal catalytic rate of AK is obtained at or near the expected physiological concentration of AMP (~300 µM (39–41)). Changes in AMP concentration will accordingly lead to changes in AK’s activity, which can substantially affect other processes in the cell by altering the availability of nucleotides. For example, we have recently shown that elevated AMP concentrations decrease the fitness of *E.coli* cells expressing inhibition-prone AK strains (27). Paalme *et al*. demonstrated that AMP levels in *E.coli* cells depend on which carbon sources are available; for example, AMP levels are elevated when only acetate is present as a carbon source (42). For mitochondrial AK it was shown that phosphorylation of AMP to ADP is inhibited when AMP levels sharply increase (for example during ischemia (43)), thus reducing the amount of adenine nucleotides available for oxidative phosphorylation (44). The current work, based on an extensive library of AK mutants and subsequent informatics analyses, shows that residues favoring substrate inhibition are often well conserved, suggesting that substrate inhibition in AK has a high functional value. Our recent study of mutant *E.coli* AK strains shows the highest bacterial growth rates for AK variants with a *K_I_* similar to the WT (27); *E.coli* AK might have evolved to represent the ideal tradeoff between efficient AMP binding and thermal stability on one side and SI on the other side (27). Even single-point mutations can change this balance, as our library data clearly shows. Interestingly, SI can be modulated through mutations that span most regions of the proteins, suggesting that long-range allosteric communication is a general feature of AK.

It is important to mention that previous work on AK dynamics suggested that the LID opening rate is the rate-determining step in catalysis (14, 15). In contrast, a recent study demonstrated an enhanced tendency toward enzyme opening for several mutants with a low catalytic activity, suggesting that domain opening is not rate-limiting at least in those mutants (45). Our smFRET data in this study and the previous one (22) demonstrate that rates of domain movement (both opening and closing) are orders of magnitude faster than the enzymatic turnover, which implies that the enzyme opens and closes multiple times before the catalytic cycle is completed. Our model of the enzymatic activity coupled with conformational changes suggests that as long as these rates are substantially higher than the catalytic turnover rate (*k*_cat_), the explicit interconversion rates between the open and closed states do not determine enzyme activity. Instead, the relative population of the two states is what determines activity and inhibition.

A matter of intense debate in enzyme catalysis is whether conformational fluctuations play a role in catalysis (2, 3, 46, 47). While several studies have claimed that enzyme fluctuations play a direct role in the chemical step, theoretical studies have refuted those (48). Our work suggests that the ultrafast domain movements help to tune the equilibrium between states of the enzyme with different activities continuously. Such a mechanism is not unique to AK. For example, we recently discovered that the activity of the AAA+ disaggregation machine ClpB is tightly correlated to the distribution between active and inactive conformational states (49). However, the interconversion between these two states happens on a much faster timescale than any other protein activity (49). Schanda et al. found that the enzymatic activity of the TET2 aminopeptidase is controlled by the conformational equilibrium of a highly dynamic loop in the catalytic chamber (50). Also, the turnover in the HisFH enzyme complex is dictated by the population of the active state, with the conversion between ground and active state taking place on a faster time scale than enzymatic activity (51).

This study goes beyond previous attempts to connect conformational dynamics and chemical steps in enzymes by developing a kinetic model that explicitly takes conformational changes into account. The model builds on the recent *in silico* observation that fast opening and closing cycles might help to find the correct mutual substrate orientation efficiently and to guarantee the high catalytic activity of the enzyme (23). It is further based on the hypothesis that *the order of binding of the two substrates to the enzyme* dictates how effectively repeated opening and closing cycles help to reorientate the substrates for the reaction. Strikingly, this model is able to correctly reproduce trends of substrate inhibition among the WT enzyme and several mutants. In conclusion, our study of a common and widely studied enzyme uncovers the mechanistic relationship between very fast conformational variation and catalysis.

## METHODS

See Supporting Information for a detailed description of the methodology for sample preparation, data collection and analysis. Briefly, a large library of AK mutants was screened for the strength of substrate inhibition. 86 unique mutants were purified and tested for their activity. Resulting inhibition constants were correlated with the conservation indices of mutated residues. Proteins for smFRET experiments were expressed and labeled with two fluorescent dyes. Single-molecule data was collected on freely diffusing molecules and analyzed using the statistical analysis algorithm H^2^MM, to obtain domain opening and closing rates. These rates served as input for simulation of enzyme kinetic curves using a model that included conformational dynamics.

## ASSOCIATED CONTENT

### Supporting Information

METHODS section containing a detailed description of the methodology, Figures S1–S11, Tables S1–S10, supporting references.

## ACKNOWLEDGMENTS

The work of G.H. was supported by a grant from the European Research Council (ERC) under the European Union’s Horizon 2020 Research and Innovation Program (grant agreement No 742637, SMALLOSTERY) and a grant from the Israel Science Foundation (no. 1250/19). The work of D.S. was supported by Deutsche Forschungsgemeinschaft (DFG, German Research Foundation, Projektnummer 490757872). The work of E.G. was supported by the National Institute of Health (NIH) grant 5R35GM139571.

## METHODS

### The degenerate library of AK

AK’s structure can be divided into three main domains – the ATP-binding LID domain (residues 118-160), the AMP-binding NMP domain (residues 30-67), and the rest, i.e., the CORE domain. The sites for mutations were chosen based on the following criteria: a) the sites should not be contacting any of the active site residues. The active site residues were defined as those residues that show >5 Å^2^ change in solvent-accessible surface area with and without the inhibitor, Ap_5_A (PDB identifier 1AKE, (1)). b) The sites should be at least 8 Å away from Ap_5_A. c) The side-chain of the WT residue should not be involved in any salt bridges or hydrogen bonds with other protein residues. 71 residues satisfied all these criteria, of which 55 were in the CORE, 10 in the LID, and 6 in the NMP domain. A degenerate codon “DHK” [D = A or T or G][H = A or T or C][K = T or G] was used to generate 13 different amino acid mutations at each of these selected 71 sites. The DHK library at each site represents all amino acid types (AILMV+DE+FY+K+NST + 1 stop codon) and hence offers a convenient alternative to “NNK” – the fully degenerate codon that codes for all 20 amino acid diversity. The mutational library at each position was generated separately by amplifying the whole plasmid (pET28a(+) with AK gene cloned between *Nde*I and *Xho*I) with the mutagenic primers. The PCR products were pooled together and transformed in *E. coli* DH5α as a single library. A single plasmid prep was done and transformed in the expression host *E. coli* BL21(DE3). A total of 1248 colonies were picked up for further assessment.

### Activity assay from cell lysate

All 1248 library candidates were grown overnight in LB in thirteen 96-well plates. This culture was diluted 1:100 in 500 μl of LB in 2 ml deep-well plates. The cultures were induced with 1 mM IPTG at OD600~0.4 (~3h) and further grown overnight before they were pelleted. The cell lysis was done using BugBuster/Benzonase mix per the manufacturer’s instructions. The cell lysate was diluted 1:4000 in 10 mM potassium phosphate buffer, pH 7.2, and used for enzyme activity as described previously (2). A quick assessment of the inhibition status was possible from the ratio of *v_0_* at 100 and 500 µM of AMP (ATP was fixed at 1000 μM) as shown in Fig. S1b.

### High-throughput protein purification

A total of 192 candidate proteins were purified – 64 each from the right, left and the middle of the distribution of 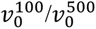. The proteins were expressed in 500 µl LB and lysed as described earlier. The his-tagged protein from the lysate was purified using HisPur™ Ni-NTA spin plate (ThermoFisher, cat # 88230). Imidazole was removed from the eluted protein solution using Zeba™ Spin desalting plate (7K MWCO) (ThermoFisher, cat # 89808), and the proteins were desalted into a 10 mM potassium phosphate buffer, pH 7.2. A first pass at Sanger’s sequencing confirmed 86 unique mutants with reliable chromatograms. Enzymatic activity assays were carried out as described previously (2). Values for the Michaelis constant (*K_M_*) and an effective inhibition constant (*K_I_*) were obtained by a fit to

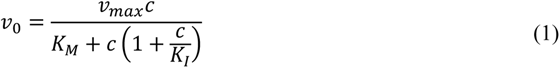

where *c* is the concentration of AMP and *ν_max_* the maximum turnover.

Subsequently, we compared the strength of inhibition (i.e. the size of *K_I_*) to the conservation index of the respective mutated amino acid. High conservation refers to a high frequency of occurrence in a multiple-sequence alignment (3, 4)). Using a Student t-test, we compared locations of mutations that led to a loss of inhibition (*K_I_* >=1500) to locations that led to an increase of inhibition (*K_I_* <=500). Positions where both inhibited and uninhibited mutations were observed (magenta in Table S1) were not considered in this analysis.

### Expression and labeling of proteins for fluorescence experiments

For the smFRET experiments, we chose proteins from the group above that demonstrated a similar or slightly higher stability than the WT (2, 5). A pET15b expression vector containing the E. coli AK C77S gene with a six-residue histidine tag at the N terminus was used to express the protein for fluorescence experiments. Alanine-to-cysteine and valine-to-cysteine substitutions were introduced at positions 73 and 142 of the protein, respectively, by site-directed mutagenesis. The sequence of AK C77S/V142C/A73C was verified by DNA sequencing. The mutant was overexpressed by transformation into E. coli BL21 competent cells. The cells were lysed using high-energy sonication or a French cell press. The pellet was then separated from the protein-containing supernatant using high-speed centrifugation. The supernatant was separated on a Ni Sepharose column (GE Healthcare HisTrap HP). Fractions containing AK were pooled and run on a second gel filtration column (HiLoad 16/60 Superdex 75 prep grade; GE Healthcare) and eluted as a single peak. Labeling reactions were performed by first incubating protein samples with Alexa 594 maleimide (Invitrogen) at a molar ratio of 75%, separating labeled from unlabeled protein on a mono-Q 5/50 GL column (GE Healthcare), and later-on labeling with an excess of Alexa 488 maleimide (Invitrogen).

### Enzymatic activity measurements of labeled protein

The enzymatic activity for the purified and the double-labeled protein variants was conducted at pH 8.0, mimicking the conditions of the single-molecule experiments. The velocity was monitored for the forward reaction (MgATP+AMP → ADP +MgADP) by the oxidation of NADH at 340 nm in coupling with pyruvate kinase and lactate dehydrogenase. The final assay mixture was: 4 nM AK, 50 mM Tris-HCl (pH 8.0), 100 mM KCl, 4 mM phosphoenolpyruvate, 5.0 mM MgCl_2_, 0.2 mM NADH, 10 units/mL pyruvate kinase, 15 units/mL lactate dehydrogenase, 0.25 mg/mL bovine serum albumin, 1 mM ATP and varying concentrations of AMP. The initial velocity was obtained by linearly fitting the NADH signal as a function of time. For the forward direction, the initial velocity of the reaction *ν_0_* was fitted to the model for two kinetically distinct pathways (6, 7):

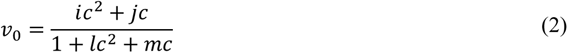

where *c* is the concentration of AMP *i*, *j*, *l*, and *m* are functions of the ATP concentration (6) and the various steps shown in Fig. 5a.

### Structure determination and refinement of AK L107I in complex with Ap_5_A

Crystals of AK L107I in complex with Ap_5_A were obtained using the sitting-drop vapor-diffusion method with a Mosquito robot (TTP LabTech). The crystals were grown from 7.5% PEG 3350, 7.5% PEG 4000, 7.5% PEG 2000, 7.5% PEG 5000 monomethyl ether, 0.07M ammonium nitrate, 2.5% ethylene glycol and 0.05M MES pH=7. The crystals formed in the space group *P22_1_2_1_*, with six copies per asymmetric unit. A complete dataset to 2.05Å resolution was collected at 100 K from a single crystal on an in-house liquid-metal-jet (LMJ) X-ray diffractometer.

Diffraction images were indexed and integrated using the CrysAlis Pro software from Rigaku, and the integrated reflections were scaled using the SCALA program (8). Structure factor amplitudes were calculated using TRUNCATE (9) from the CCP4 program suite. The structure was solved by molecular replacement with the program PHASER (10), using the structure of AK from *E. Coli* (PDB code 4JZK).

All steps of atomic refinement of both structures were carried out with the CCP4/REFMAC5 program (11) and by Phenix refine (12). The models were built into *2mF_obs_ − DF_calc_*, and *mF_obs_ − DF_calc_ maps* by using the COOT program (13). Details of the refinement statistics of the AK L107I mutant in complex with Ap_5_A structure are provided in Table S10. The coordinates were deposited in the RCSB Protein Data Bank (PDB:8BQF).

### smFRET data acquisition

Single-molecule data was acquired on freely diffusing molecules using a Microtime 200 system (PicoQuant). Flow cells were prepared as described previously (14) and filled with a mixture of 15 pM labeled enzyme, 50 mM Tris-HCl (pH 8.0), 100 mM KCl, 5 mM MgCl_2_, 0.01% Tween (Thermo Fisher), and substrates (ATP, ADP, AMP; Sigma). Importantly, we verified that our ATP solutions did not contain any ADP using ^31^P-NMR spectroscopy. Substrate concentrations used for experiments under turnover conditions are given in Table S5. The appropriate ADP concentration to guarantee equilibrium (zero flux) was calculated using the following rate equation:

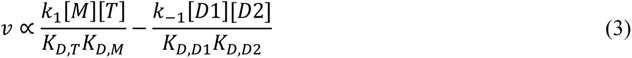

where k_1_/ k_-1_ are the forward and backward rate constants for the reaction and [M] and [T] are the AMP and ATP concentrations. K_D,S_ is defined as the dissociation constant of the respective substrate (T, M, D1, D2) and were taken from Sheng *et al.* (15). [D_1_] and [D_2_] are the concentrations of ATP bound to the LID and NMP domain, respectively. [D_2_] is calculated using

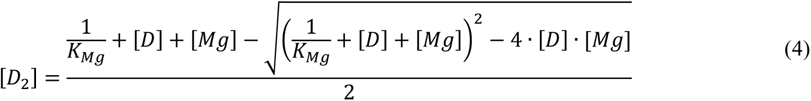

where [Mg] and K_Mg_ (15, 16) are the concentration and dissociation constant for magnesium, respectively. FRET efficiency histograms from the first and last 1h of each measurement were shown to overlap, validating that indeed the substrate concentration did not change during the measurement.

Measurements were conducted in the pulse-interleaved excitation mode, using a sequence of one pulse for acceptor excitation (594 nm, power 10 μW) and three pulses for donor excitation (488 nm, power 50 μW) at 40 MHz. The laser beams were focused 10 µm deep into the sample solution. Molecules diffusing through the beam emitted short bursts of photons that were divided into two channels according to their wavelengths, using a dichroic mirror (zt594rdc; Chroma) and filtered by band-pass filters (HC520/35 (Semrock) for the donor channel and ET 674/75m (Chroma) for the acceptor channel). Arrival times of these photons were registered by two single-photon avalanche photodiodes (SPCM-AQRH-14-TR, Excelitas) coupled to a time-correlated single-photon counting module (PicoHarp 400, PicoQuant).

### Photon burst selection

Fluorescent bursts in the single-molecule data were detected using methods developed in the lab (14, 17, 18). A cut-off time of 5 μs between individual photons was determined from the histogram of the time lags and used to find the effective start and end points of each burst. Only fluorescent bursts with a total of 50 photons or more were selected for further analysis, resulting in an average photon flux of around 400 photons per millisecond. The raw FRET efficiency of each burst was calculated based on the photons detected in both channels following donor excitation only. The raw stoichiometry was obtained from the detected photons in both channels after both excitations, as described elsewhere (19, 20). A 2D histogram of raw stoichiometry versus raw FRET efficiency was generated, from which we extracted the amount of emitted donor photons leaking into the acceptor channel and the level of direct excitation of the acceptor dye by the 485 nm laser. The photon stream in both channels was corrected for the leakage of photons from the donor to acceptor, using the apparent FRET efficiency of donor-only molecules (19, 21). The correction factor for direct excitation of the acceptor dye was determined from the acceptor-only population (19, 21). The FRET efficiency for each burst was calculated from the corrected number of photons arriving from the acceptor channel divided by the total number of photons. To obtain the final FRET histogram without the donor-only and acceptor-only populations, we selected only photon bursts with a stoichiometry corresponding to molecules with both donor and acceptor dyes.

### Analysis of protein dynamics with H^2^MM

To extract the dynamics hidden in the photon bursts, we used the H^2^MM algorithm (17). Only double-labeled molecules and photons arising from donor excitation were taken for this analysis. For each protein variant, the FRET efficiencies of the open and closed state were optimized globally to give the best fit across varying substrate concentrations. This procedure was employed since the structures of the two states themselves were considered to be unaltered by substrate binding, in contrast to the distribution between the states. On the other hand, the initial populations of the states and the interconversion rates were optimized for each measurement separately. Similar FRET efficiency values were obtained for all mutants, 0.37±0.02 / 0.72± 0.02 for the WT and L107I, 0.38±0.01 / 0.68±0.02 for L82V and 0.40±0.01 / 0.70±0.02 for F86W for the open and closed state, respectively. This reflects the fact that the mutations do not significantly alter the overall protein structure. In the case of L107I, this could be additionally verified by the excellent agreement of the crystal structure of the closed state reported in this work with that of the WT (1).

For each mutant, at least two different data sets (independently prepared samples) were analyzed for all different substrate concentrations. The resulting parameters were validated with different tests: A recoloring analysis (Fig. S5), a visualization of the impact of the interconversion rate on the FRET efficiency histogram (Fig. S6), a dwell-time analysis (Fig. S7, Table S3) and fluorescence correlation spectroscopy experiments (Fig. S8, Table S4). Details are given in the paragraphs below. Representative single-molecule trajectories are shown in Fig. S4. These include the assignment of the most likely state-sequence using the Viterbi algorithm on a photon-by-photon basis (17).

Our kinetic model does not take into account dye blinking, as blinking events are largely filtered out in our rigorous burst selection. Indeed, less than 0.1% of our selected trajectories show a maximum interphoton times of more than 100 μs after acceptor excitation. Blinking of the acceptor dye would manifest as gaps in this photon stream. As a test, we have also carried out H^2^MM analysis on data sets where bursts with a potential blinked state (Δt>100 µs after acceptor excitation) had been filtered out, and saw minimal differences compared to the complete data sets. For a visualization of the photon streams after both donor and acceptor excitation, see Fig. S4.

### Recoloring analysis

We performed a recoloring analysis to verify the parameters obtained from the H^2^MM analysis (14, 22). In this method, the arrival times of photons in each data set are retained, but their “colors” (i.e. whether they belong to the donor or acceptor) are erased. A stochastic simulation based on the H^2^MM parameters is then used to reassign the photons to the two experimental channels, and FRET efficiency histograms are reconstructed. A good match between the original histograms and the recolored ones indicates a successful analysis.

### Dwell-time analysis

The dwell-time analysis yields the distributions of times that the protein spends in each state (in this case, the open and closed state). Here we computed these he distributions using a likelihood-weighted segmentation algorithm developed in-house (17, 23). In this analysis, for each burst every possible sequence of states contributes to each dwell time a fraction of a count, equal to the likelihood of the sequence. In contrast with the more common dwell-time analysis based on the Viterbi algorithm, here all possible state sequences are taken into account, not only the most likely one. A good agreement of rates obtained directly from the H^2^MM analysis and those obtained from the dwell-time analysis was considered as a validation of the analysis (Table S3).

### Fluorescence correlation spectroscopy

FCS experiments were performed on freely diffusing molecules using the Microtime 200 microscope (PicoQuant) at a power of 20 μW (488 nm), at a concentration of 1 nM of the double-labeled WT protein and varying concentrations of ATP. For the cross-correlation between donor and acceptor photons, identical optical elements to those mentioned in the chapter “smFRET data acquisition” were used. Auto-correlation between donor photons was obtained by first filtering the light using a band-pass filter (HC520/35 (Semrock) and then splitting it between two detectors (to reduce the effect of afterpulsing). Correlation functions were calculated using the SymPhoTime 64 software (PicoQuant). To isolate the conformational dynamics from the diffusion contribution to the correlation functions, for each sample we calculated the ratio between the cross-correlation of donor and acceptor (*CC_DA_*) and the auto-correlation of donor photons (*AC_DD_*), based on the procedure of Torres and Levitus (24):

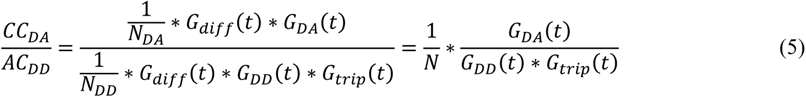

This operation cancels out the diffusion term *G_diff_ (t)* in *CC_DA_* and *AC_DD_*. *N_DA_*,*N_DD_* describe the number of molecules containing both labels and the donor label, respectively. These numbers differ due to the presence of donor-only labeled molecules and photobleaching (25). We combine both terms into an effective amplitude *N*. The remaining terms attribute for conformational (*G_DA_ (t), G_DD_ (t)*) as well as triplet dynamics (*G_trip_ (t)*). Once the diffusion term is cancelled, we use the following approximate equation for fitting:

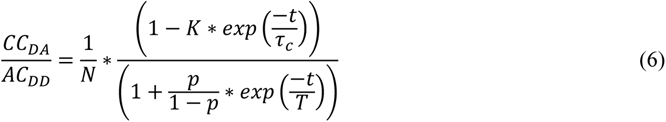

*τ_c_* is the time constant for conformational dynamics. The pre-exponential factor K depends on kinetic rates and the relative visibility of each state (24). *T* is the triplet relaxation time and *p* the fraction of dye molecules in the triplet state. *T* and *p* are independent of substrate concentration and were optimized globally.

### Differential scanning fluorimetry

Temperature melts of AK variants were carried out using a BioRad CFX384 real time PCR machine with the SYPRO orange dye. The final concentration of protein used was 4 μM, and the solution contained SYPRO orange at 5×, 5 mM of MgCl_2_, and varying concentration of ATP (0-20 mM) or AMP (0-40 mM). The total volume was adjusted to 10 μl with 10 mM potassium phosphate buffer, pH 7.2. The melting temperature (Tm) was estimated as the temperature at which the melt-curve derivative peaks. This method yields a proxy for binding as it essentially determines the stabilization of a protein by its ligand. The effect of mutation on ATP or AMP can be qualitatively determined from curves of Tm as a function of the ligand concentrations.

### Simulation of the enzymatic activity

Activity curves were simulated with the program KinTek Kinetic Explorer (KinTek, Snow Shoe, PA), with experimentally derived protein dynamics parameters and substrate affinities as input. Opening and closing rates were obtained by fits to the apparent rates observed in the smFRET experiments, as described in Supplementary Note 1. The obtained parameters are given in Table S8. The rate of productive binding events (*k_1_*) was derived from the enzymatic activity data, using the ratio *k*_cat_/*K*_M_ (often referred to as the specificity constant). With the Michaelis constant *K*_M_ given as

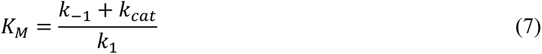

and *k*_-1_ as the rates for substrate release, *k*_cat_/*K*_M_ can be written as

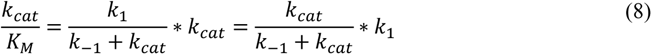

and approaches *k*_1_ when the rate of unproductive product release *k*_-1_ is minimal. The resulting parameters are given in Table S9.

**Supplementary Note 1. Substrate dependency of the closed state occupancy.** The occupancy of the closed state in AK is dependent on the substrate concentration (Figure 3). In particular, the binding of ATP stabilizes the closed conformation. To account for this behavior, we assume that each substrate-bound species in Figure 5a has a specific distribution between the open and closed conformation.

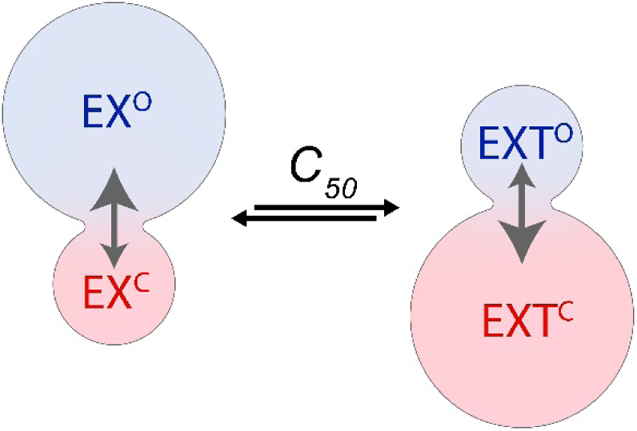

In this scheme, EX denotes the enzyme in its apo (EX=E) or AMP bound (EX=EM) form, and EXT is the enzyme already bound with ATP. The uppercase O/C indicates the protein conformation, open or closed, respectively. In the open state, the enzyme can bind or release ATP. The ratio between the different forms is given by the substrate concentration and *C_50,ATP_*, i.e. the concentration which triggers half of the maximal conformational change. Each enzyme form contributes to the apparent closed state occupancy *Occ_app_* according to its population.

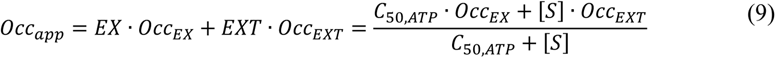

with *[S]* as the substrate concentration. *[S]* hereby refers to ATP, when it is the sole substrate, or ATP+ADP when all three substrates are present. Parameters obtained from this model are given in Table S6.

For the addition of AMP to the ATP bound form (ET), we found that minor concentrations (<1 mM AMP) did not induce changes. Therefore, our single-molecule experiments cannot distinguish between the ET state and the non-inhibited ETM state. In contrast, inhibitory concentrations of AMP above 1 mM trigger an increase in the apparent closing rate. We attribute the faster closing to the inhibited 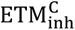 state. Similar to Eq. (9), in this scheme each enzyme form contributes to the apparent closing rate *k_c,app_* according to the population of the inhibited E_inh_ and non-inhibited E_non_ states (ET+ETM).

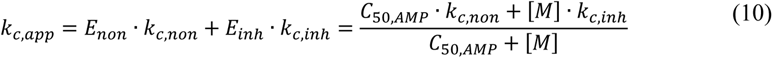

with *[M]* as the AMP concentration. Parameters obtained from this model are given in Table S7.

---

**Figure S1:**
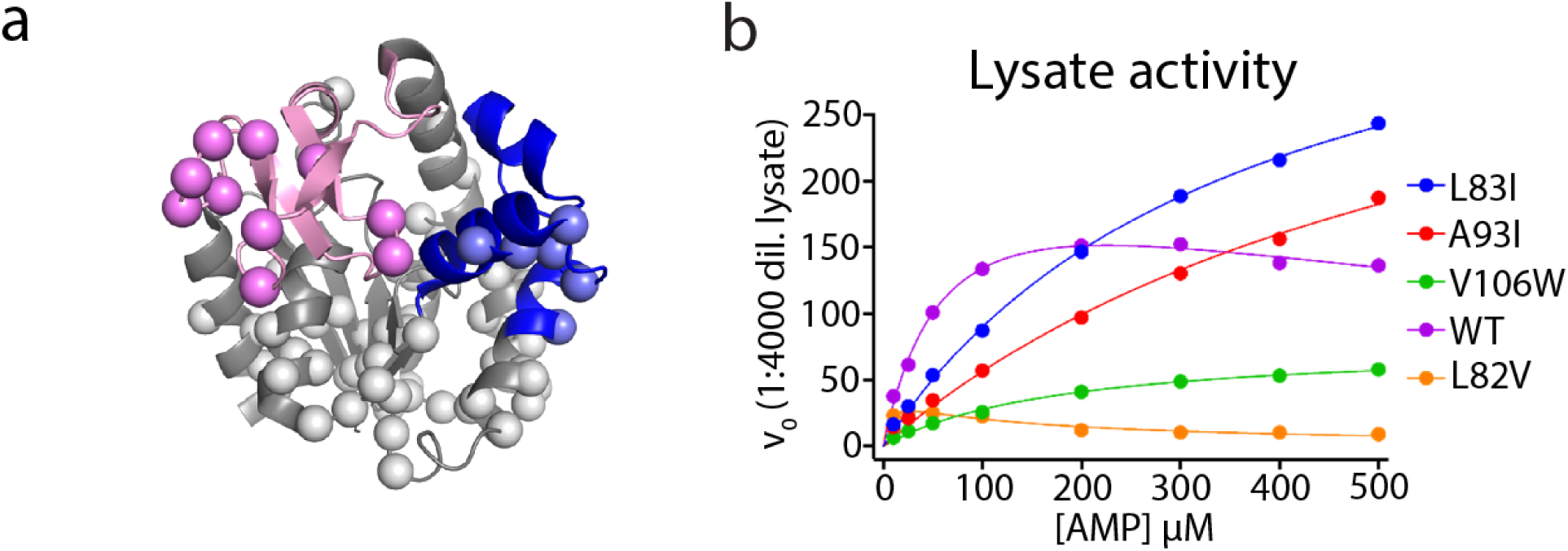
Construction and analysis of the AK mutant library. a) The domains are colored as Core in gray, LID in pink, and NMP in blue. Spheres represent positions chosen for library generation. 71 residues were selected that were at least 8 Å away from the inhibitor AP_5_A and whose sidechains were not involved in any hydrogen bonding with rest of the protein. 55 of these positions are present in the Core domain, 10 in the LID domain and 6 in the NMP domain. b) In-lysate activity measurements of known mutants of AK confirms that such measurements can reproduce the inhibition profile of the corresponding purified proteins.

**Figure S2:**
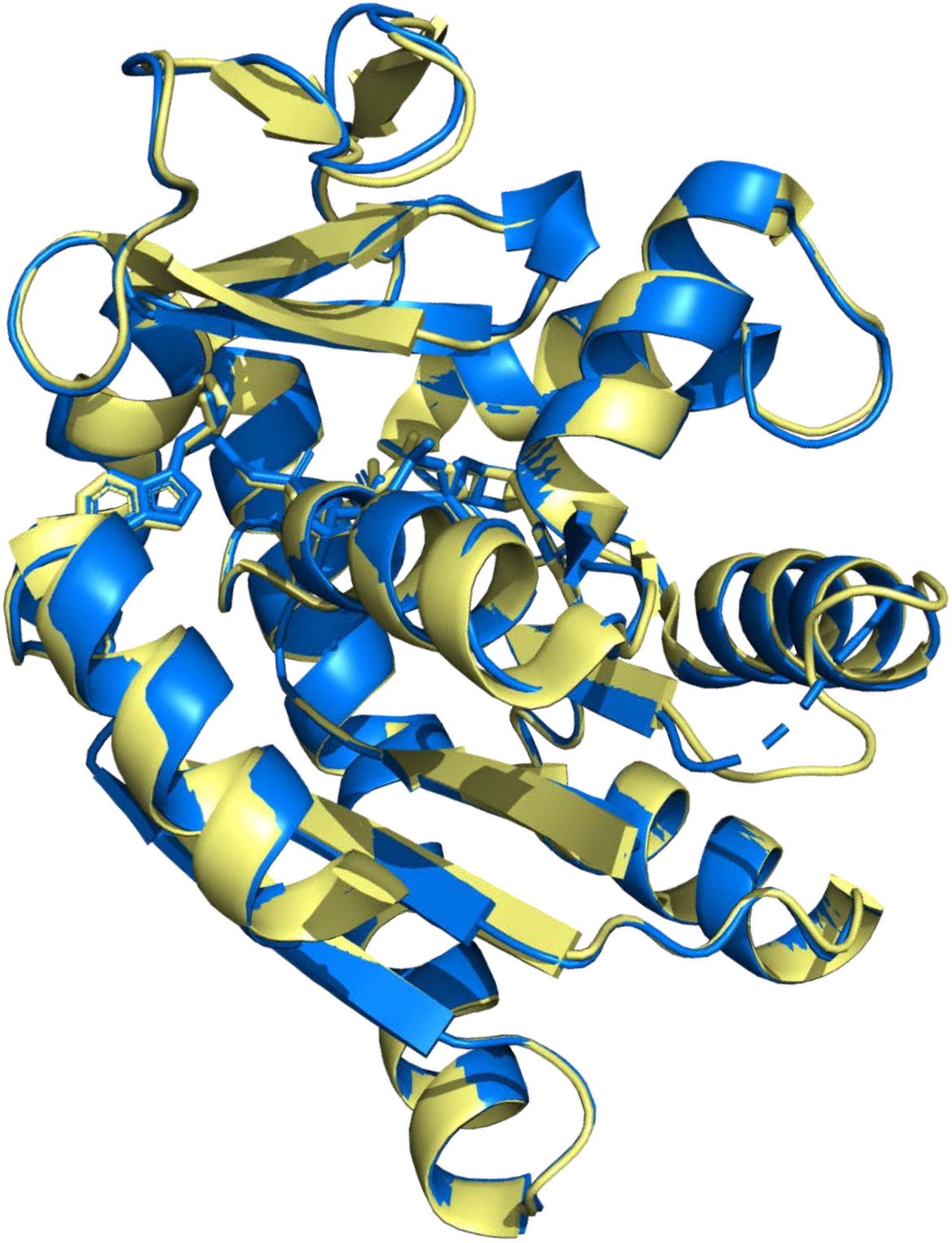
Overlay of the crystal structure of the WT in yellow (PDB:1AKE (1)) with the L107I mutant (blue). Both structures are in complex with the ligand Ap_5_A. Shown is the respective first unit of the asymmetric unit.

**Figure S3:**
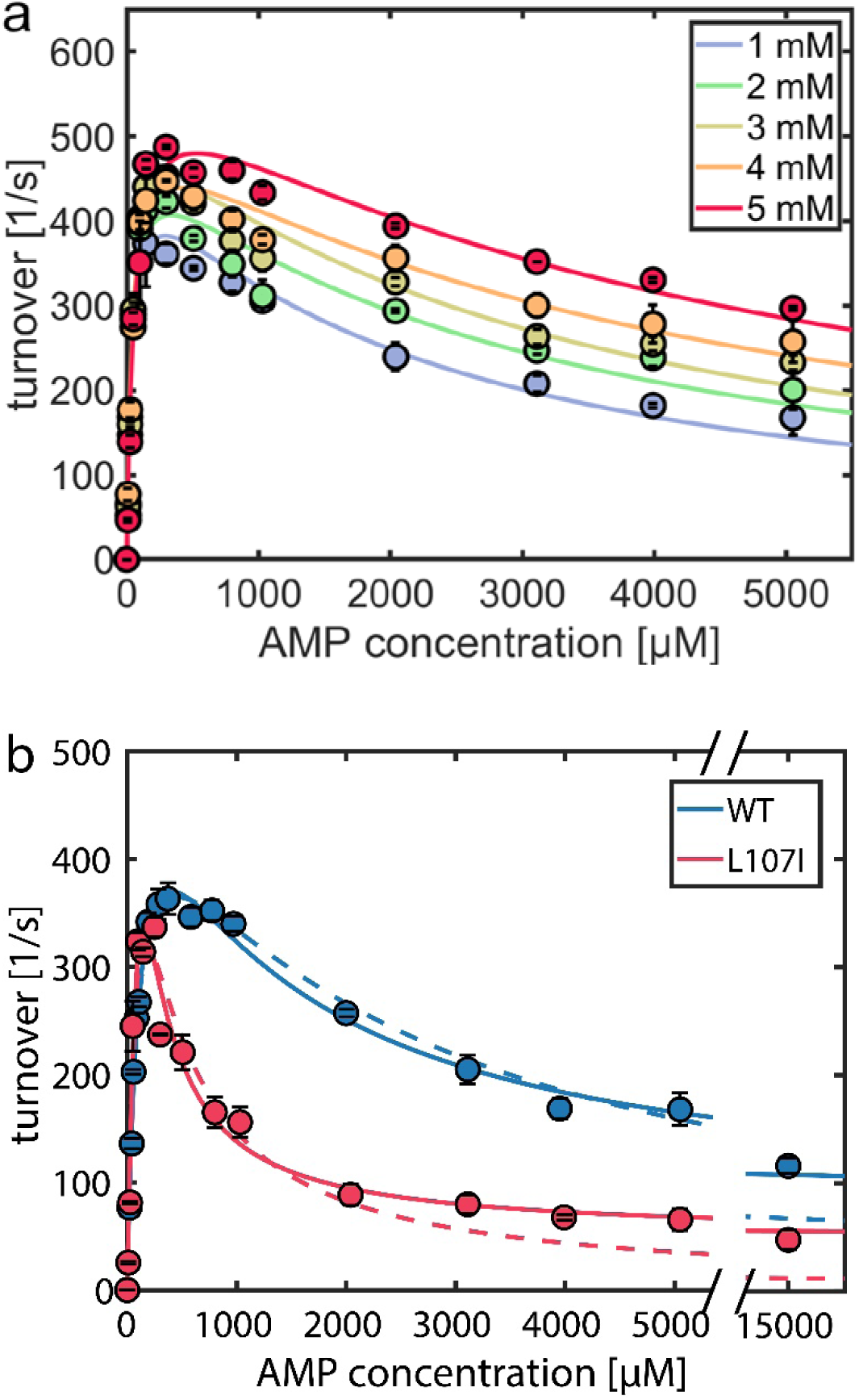
Impact of ATP and AMP on inhibition. (a) ATP lifts substrate inhibition by AMP in a dose-dependent manner for the WT. Kinetic parameters based on an inhibition mechanism involving a dead-end complex are given in Table S2. (b) Enzymatic activity for WT (blue) and L107I (red) as a function of AMP concentration. Solid curves indicate fits for a model with two alternating pathways as described (6). For comparison, fits to an inhibition mechanism with dead-end complexes are shown as dashed lines. The enzyme shows residual activity at high AMP levels, in agreement with previous studies (5, 26, 27), suggesting that the loss in activity cannot be attributed to the formation of a dead-end complex (e.g. AMP bound to the ATP site) but to a kinetically impaired pathway. At low AMP levels, only a slight deviation between the two models is seen.

**Figure S4:**
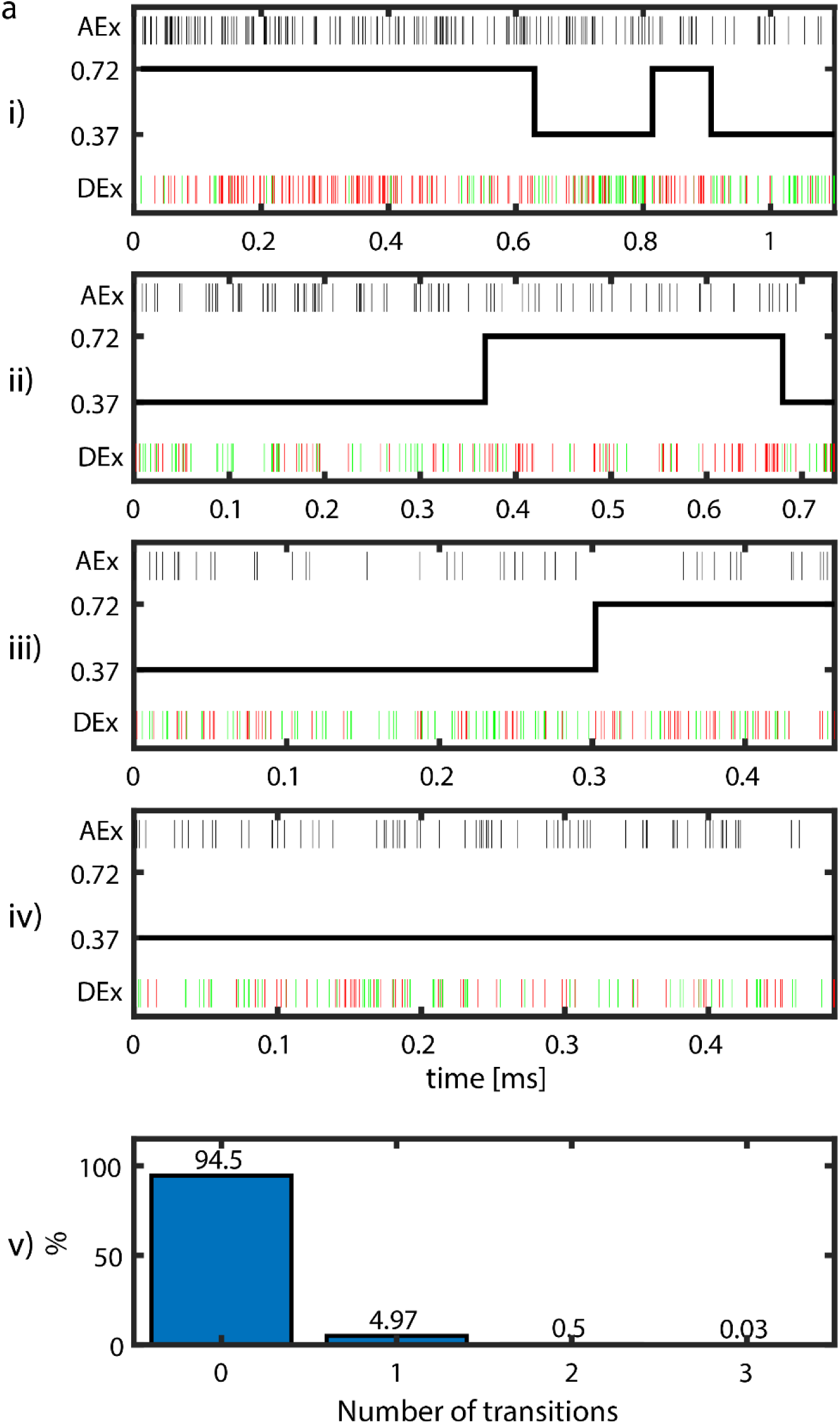
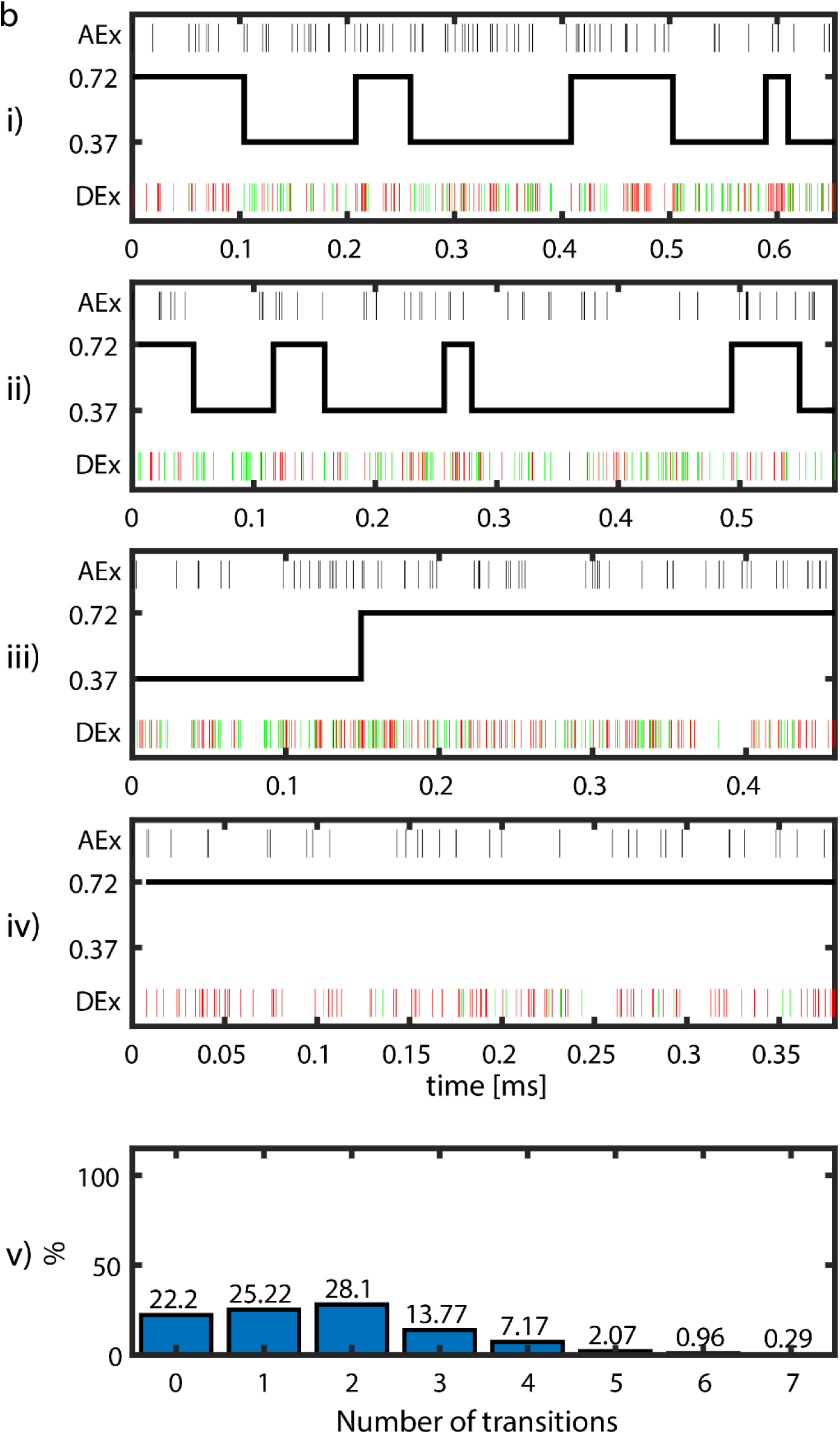
Representative photon-by-photon trajectories and Viterbi assignment. Representative single-molecule trajectories are shown for the WT protein under a) the apo condition and b) 1mM ATP. For i-iv), the top panel shows the arrival time of photons after the acceptor pulse. The bottom panel shows the arrival time of photons after the donor pulse, in green for donor photons and red for acceptor photons. The black solid line depicts the most likely state sequence according to a Viterbi assignment (17). v) depicts the distribution of the number of transitions identified by the Viterbi algorithm in each data set. Under the apo condition (a), only a few trajectories show more than one transition (i+ii). Most trajectories show either one (iii) or no transition (iv). In the presence of ATP (b), domain opening and closing are faster (Table S3), increasing the probability to observe a transition within a single trajectory.

**Figure S5:**
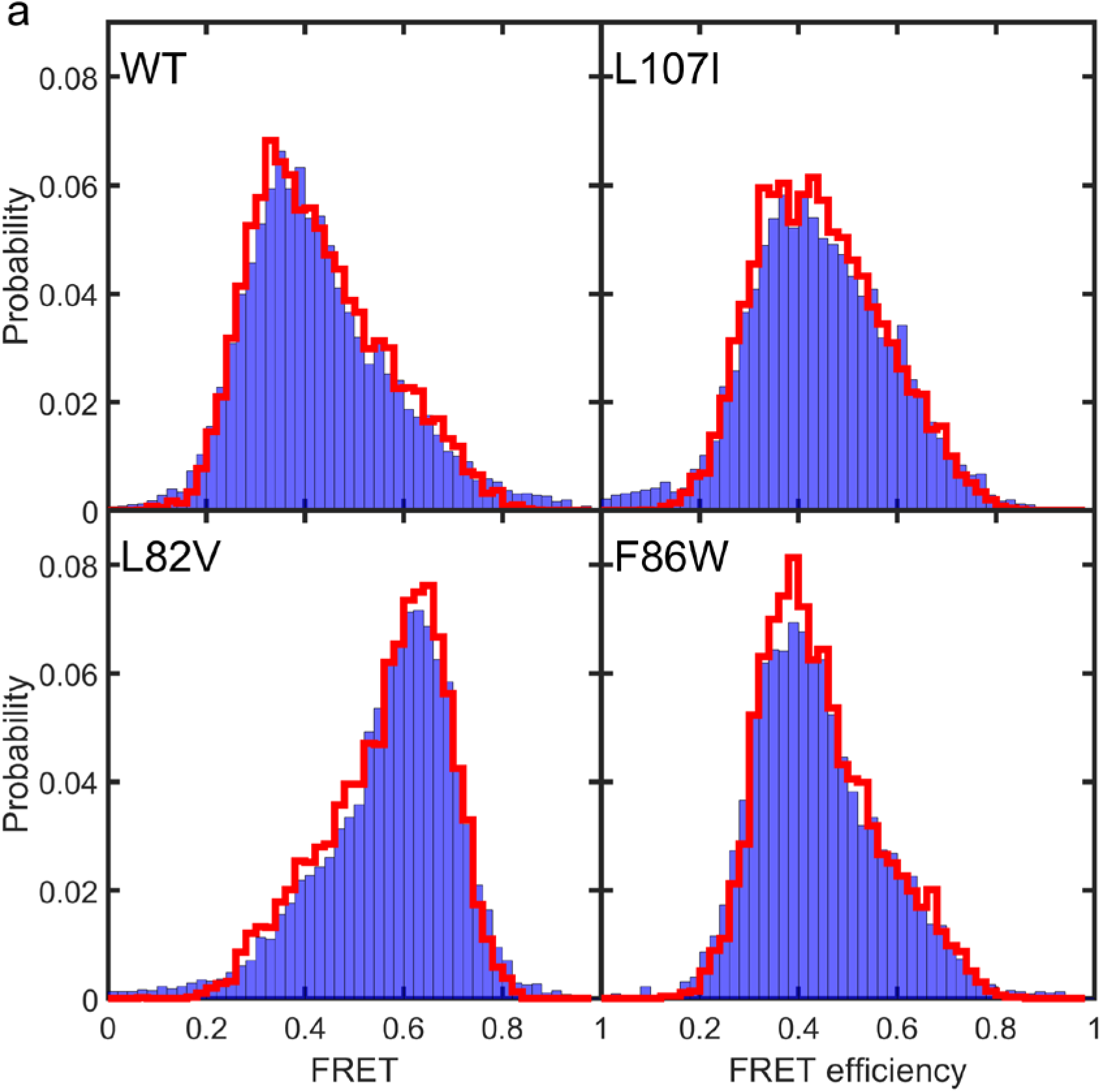
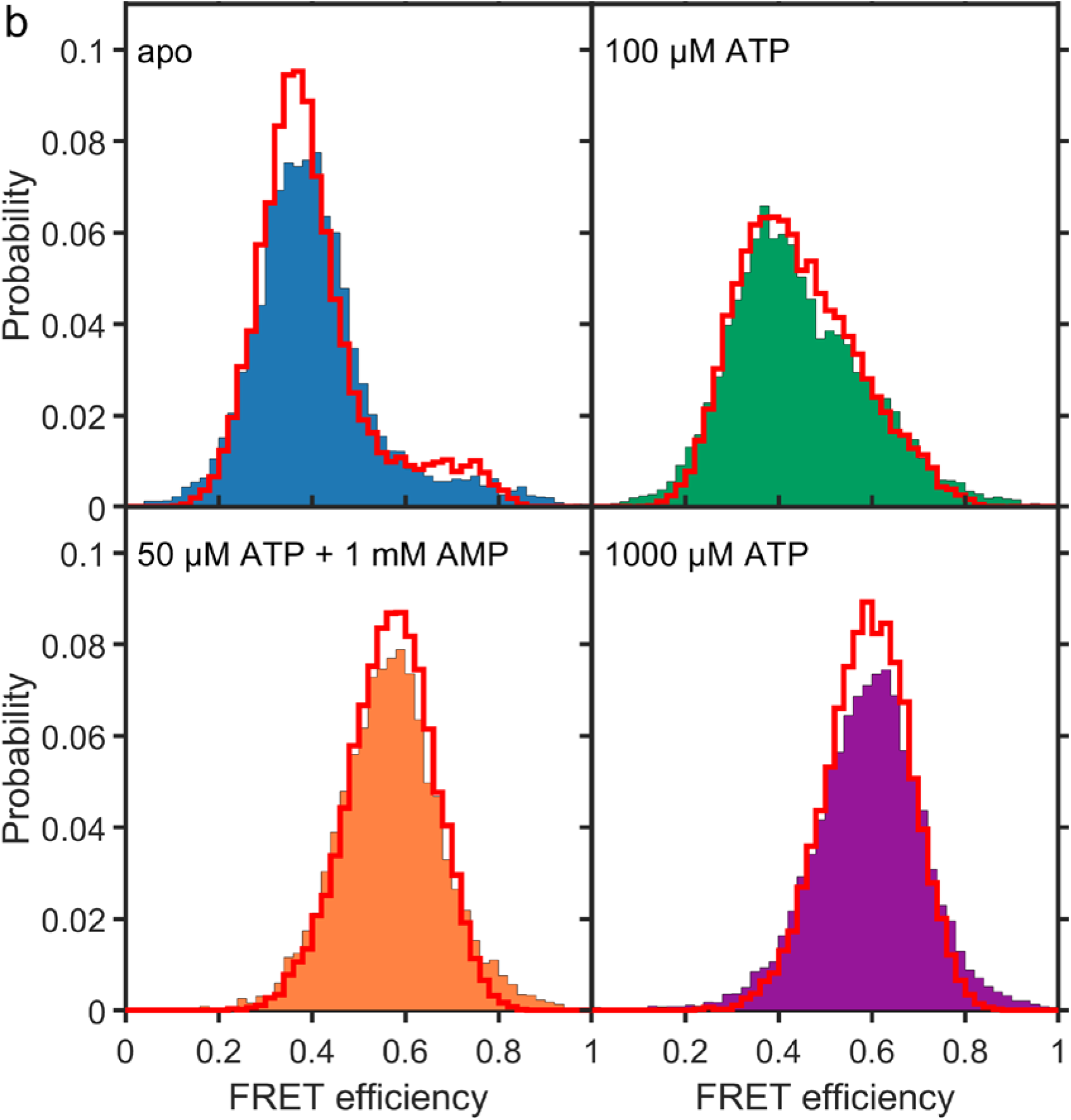
Validating H^2^MM models using recoloring. a) Representative experimental histograms for the different protein variants are shown in blue at an ATP concentration of 100 µM. The recolored histograms are depicted as solid red lines and show very good agreement between simulation and experiment. b) The model also attributes well to the effect of substrate binding. Shown are recolored histograms as red lines for the WT at the substrate concentrations depicted in Fig. 2a, in matching colors: blue for the apo protein, green for 100 µM ATP, orange for 50 µM ATP with 1 mM AMP (orange) and purple for 1000 µM ATP.

**Figure S6.**
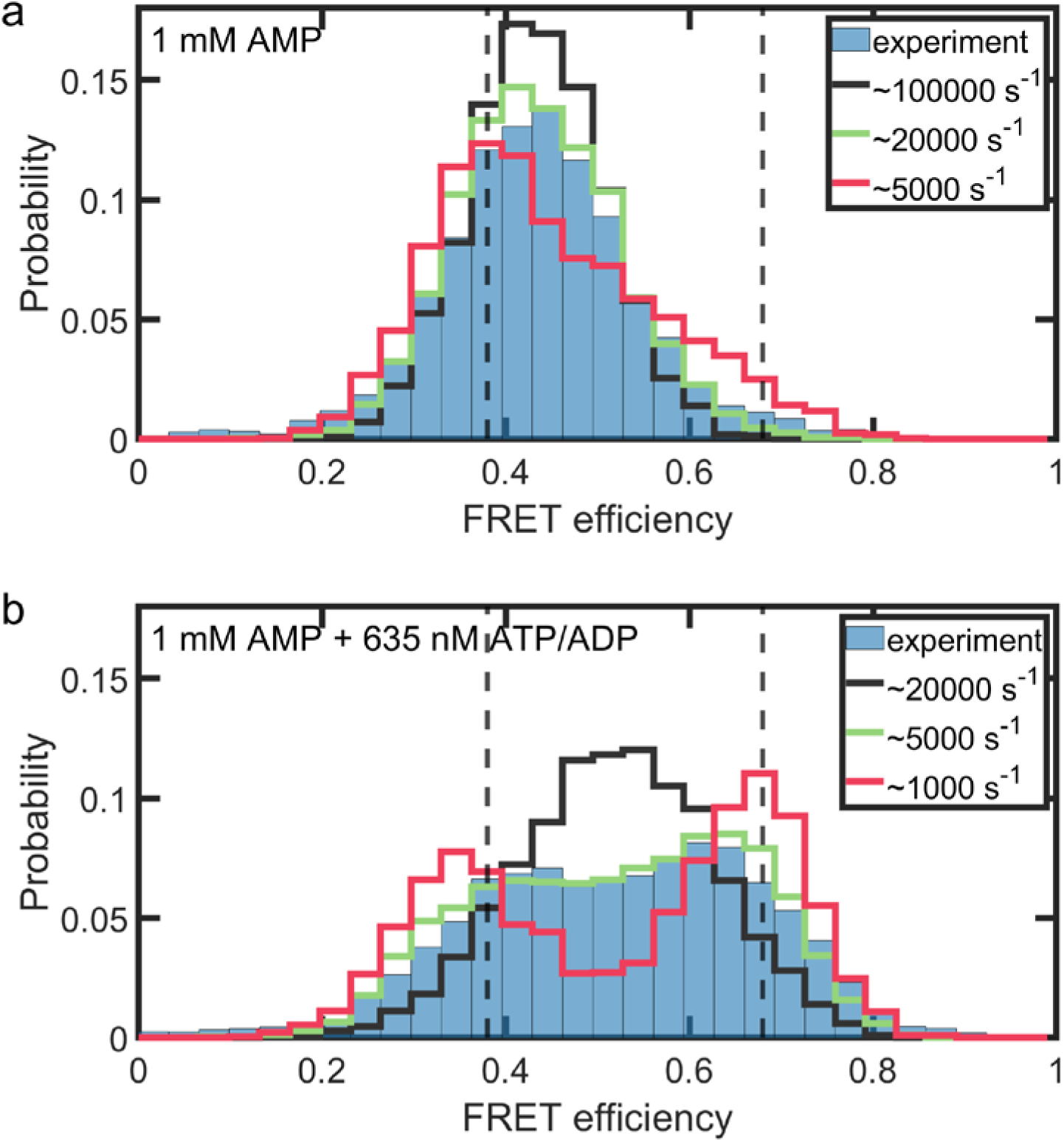
Visualizing the effect of interconversion rates on the FRET histogram. The experimental FRET histogram of L82V (blue) is plotted together with recolored histograms, reflecting the distribution of open/closed states from H^2^MM analysis but with altered interconversion rates between the states. The dashed lines indicate the most likely positions of the open (0.38) and closed state (0.68) for L82V obtained from the analysis. The green curve depicts the histogram based on the rates from the H^2^MM analysis, the black curve is for 4 times faster conversion rates, and the red curve for 4 times slower rates. (a) The protein shows a fast interconversion in the presence of AMP alone, with the conformational equilibrium in favor of the open conformation. (b) Adding a minor amount of ATP/ADP (combined concentration 625 nM) shifts the population towards the closed state (compare to Fig. 3c). Due to the slower interconversion, the histogram splits up into two peaks, as the average dwell time of the states exceeds the average burst length. In both a+b), the rates derived by H^2^MM (green) give the best fit to the experimental data.

**Figure S7.**
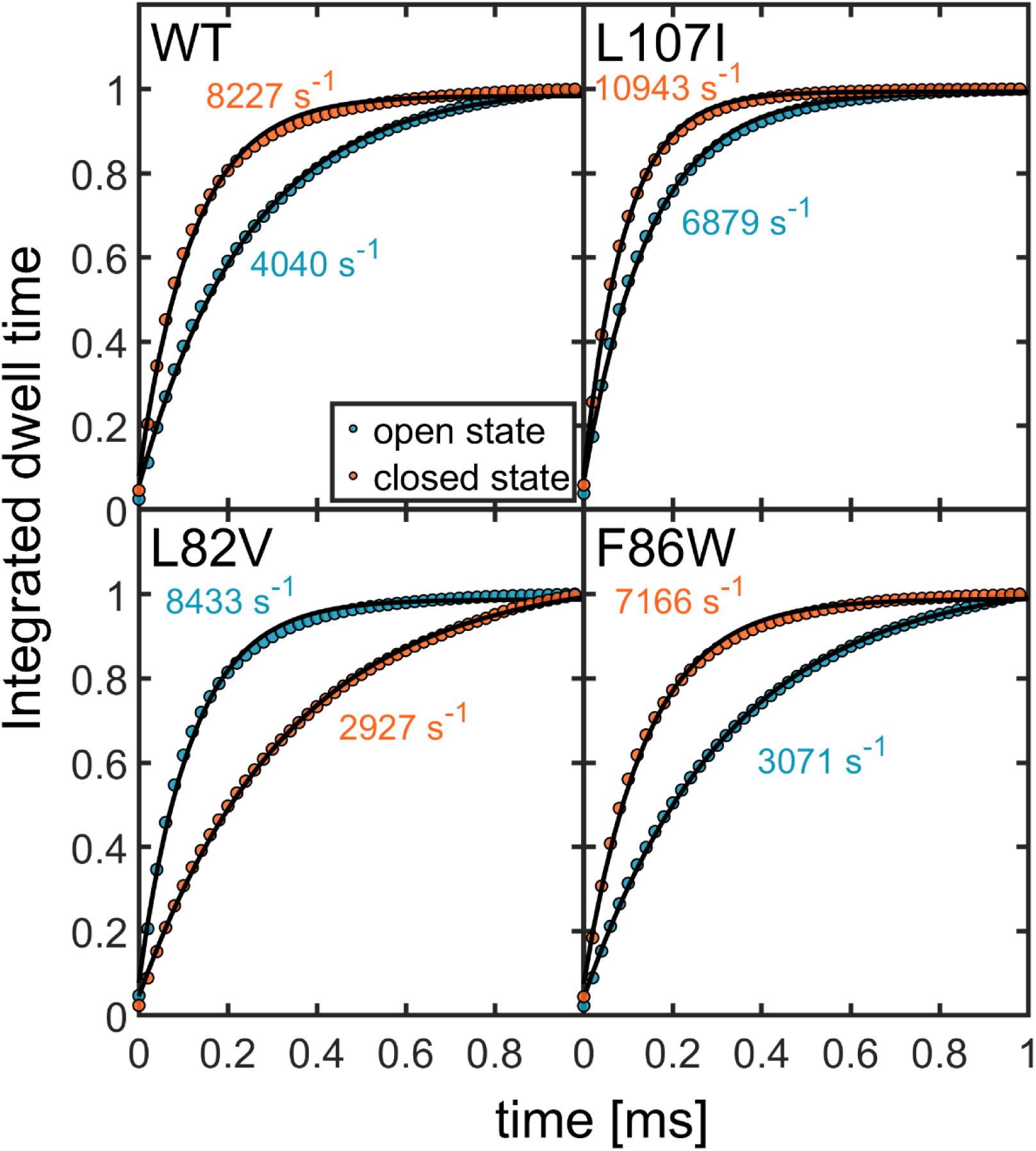
Dwell times analysis. Integrated dwell-time distributions are shown for the open state (cyan) and the closed state (orange) for the four different protein mutants as in Fig. S6a (ATP concentration 100 µM). Black lines are fits to single-exponential functions. Both closing and opening rates extracted from the dwell time distributions compare very favorably to the rates obtained by H^2^MM analysis (Table S3).

**Figure S8.**
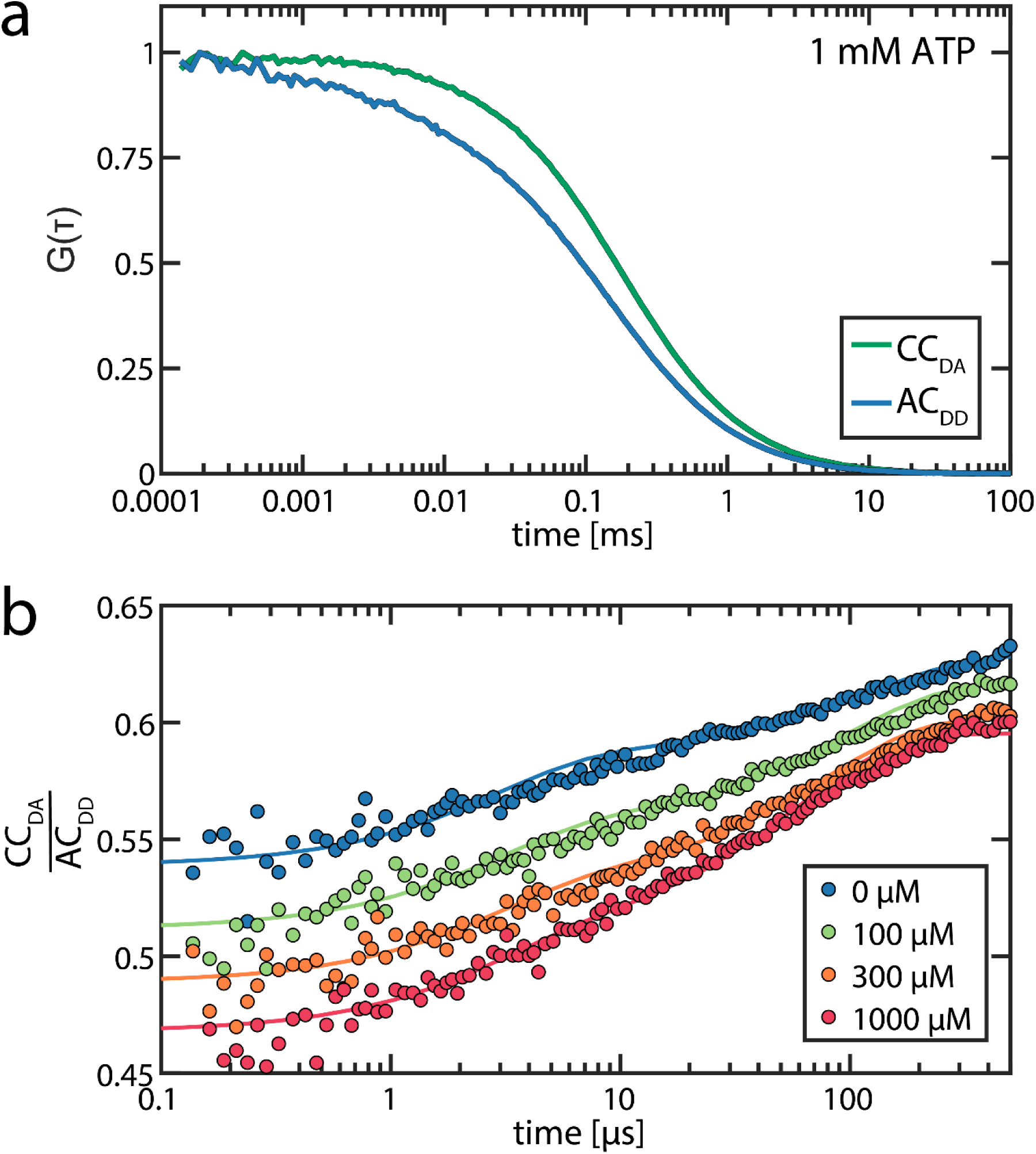
Fluorescence correlation spectroscopy. As an independent control for the presence of fast conformational dynamics in AK, we performed fluorescence correlation spectroscopy on the double-labeled WT protein. (a) Green-donor and acceptor cross-correlation (CC_DA_) at an ATP concentration of 1 mM, blue-donor auto-correlation (AC_DD_). Curves were normalized for a better comparison and show the loss of correlation due to the diffusion of molecules. In AC_DD_, correlation is lost faster due to the presence of conformational dynamics on the microsecond timescale (and triplet kinetics). b) To isolate the conformational dynamics from the diffusion part, the ratio CC_DA_/AC_DD_ is calculated (28). Shown are measurements at ATP concentrations of 0 µM (blue), 100 µM (green), 300 µM (orange) and 1000 µM (red). The curves were fit to equation 6, parameters are given in Table S4. With increasing ATP concentration, the time constant for conformational dynamics τ_c_ decreases as the conformational dynamics are getting faster, in agreement with our H^2^MM analysis. Simultaneously, the amplitude *K* increases.

**Figure S9:**
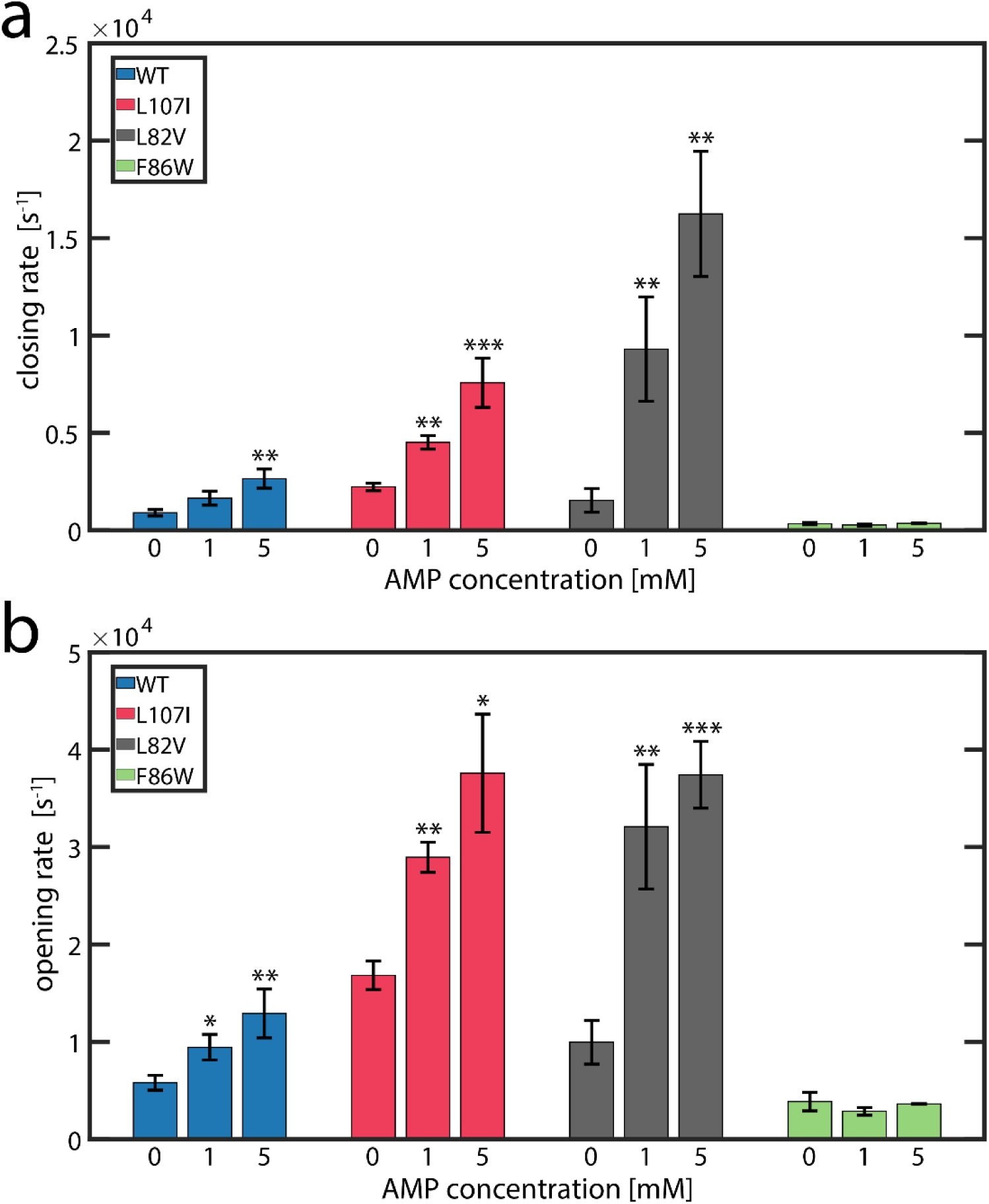
Inhibited protein variants show faster protein dynamics in presence of AMP. Closing (a) and opening (b) rates are altered for all mutants, except for the non-inhibited F86W. Asterisks indicate the significance of the deviation of parameters from the apoprotein (***: p<0.01, **: p<0.05, *: p<0.1, no index: p>0.1, t-test). Error bars are given as the standard error of the mean of 3 independently prepared samples.

**Figure S10:**
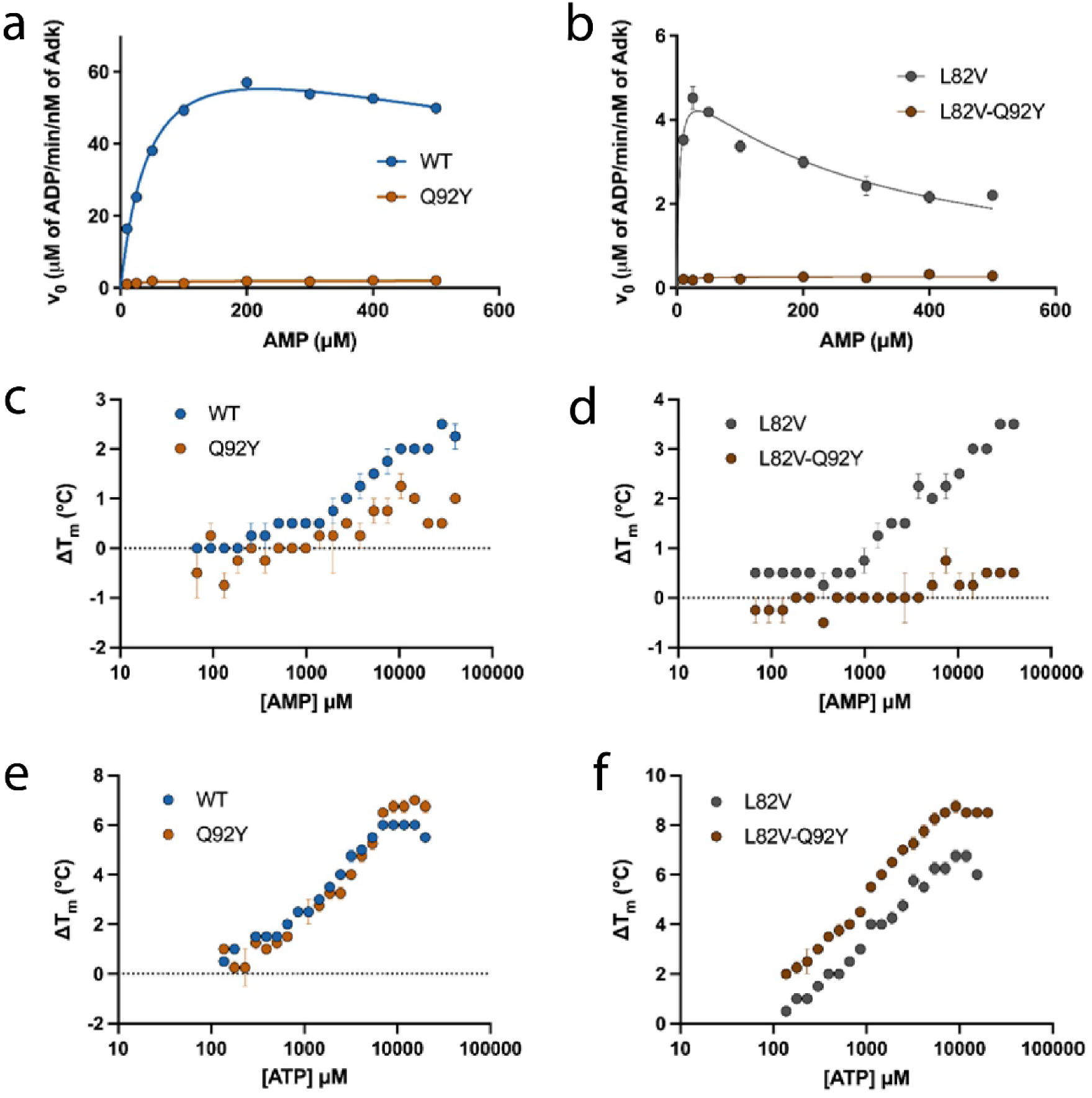
Number of AMP binding sites probed by Differential Scanning Fluorimetry. a-b) Impact of the Q92Y mutation on enzymatic activity for the WT (a) and L82V (b). The Q92Y mutation abolishes crucial interactions between the substrate and the sidechain of Q92 (1), rendering the enzyme inactive. c-d) Titration of WT/WT-Q92Y (c) and L82V/L82V-Q92Y (d) proteins with AMP using Differential Scanning Fluorimetry. The proteins were mixed with different concentrations of AMP and the thermal melting temperatures (T_m_) were monitored using the fluorescence of SYPRO Orange. ΔT_m_ refers to the change in melting temperatures with respect to the unliganded protein. The Q92Y mutation prevents AMP binding. e-f) Titration of WT/WT-Q92Y (e) and L82V/L82V-Q92Y (f) proteins with ATP. The ability to bind ATP is not affected by the Q92Y mutation.

**Figure S11:**
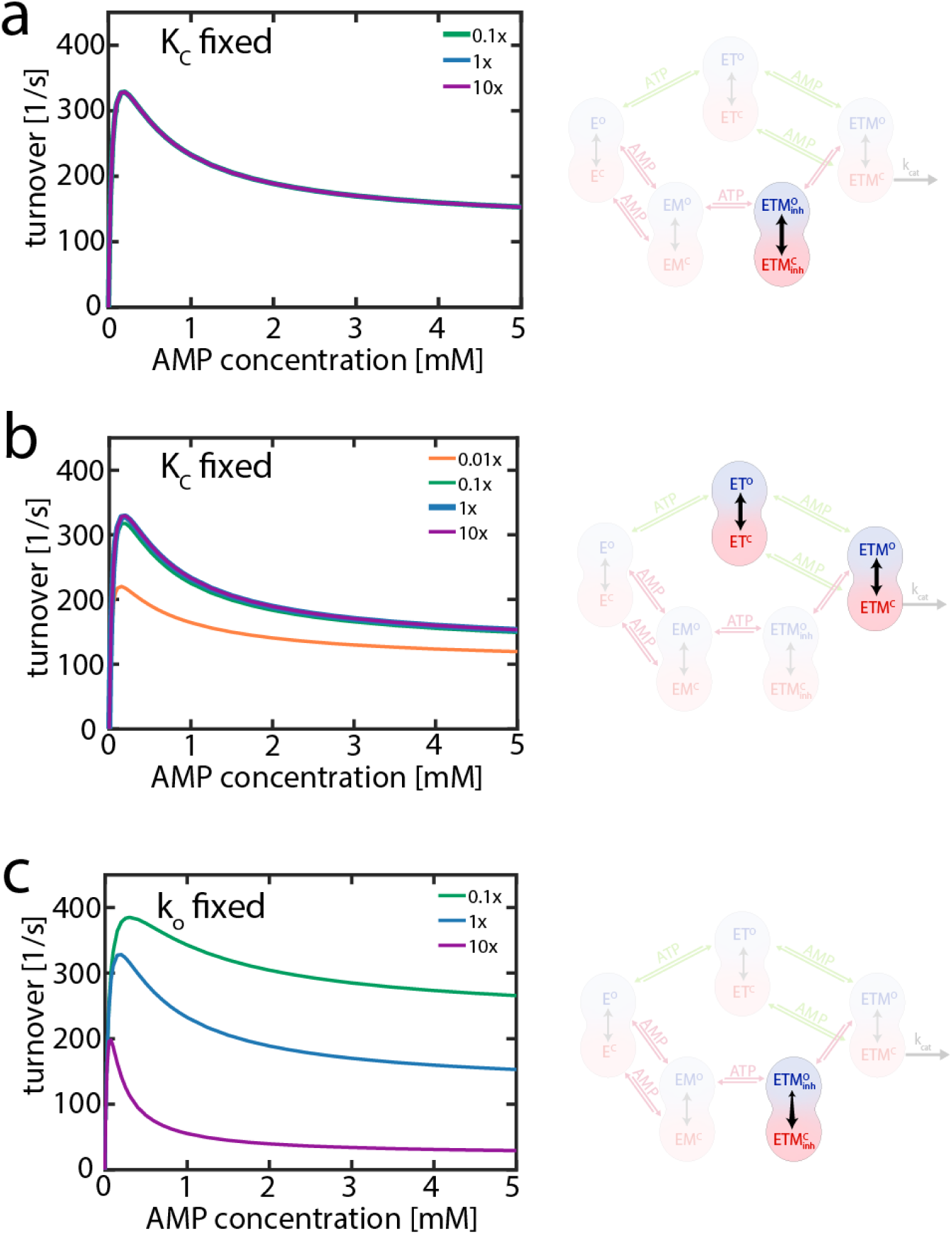
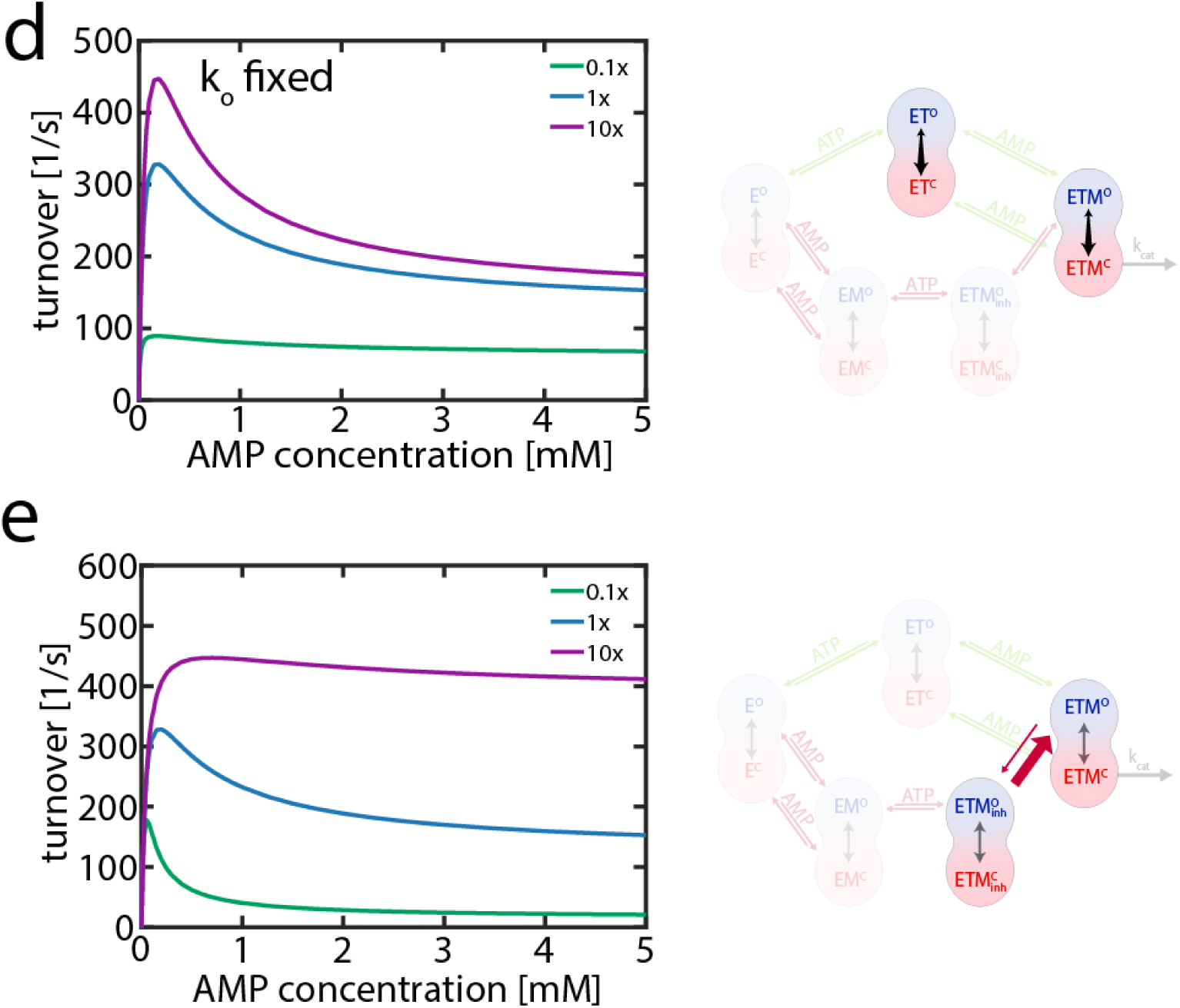
Effect of protein dynamics on simulations of enzymatic activity. The role of conformational dynamics in enzymatic activity was assessed further by altering the rates for specific processes within the conformational cycle (Figure 5). In the left panel, the experimental rates (blue) were scaled by a factor of 0.01 (orange), 0.1 (green) or 10 (purple). The right panel visualizes which rates have been altered. (a) Changing the rate constant for the unproductive closing (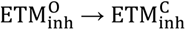) has no impact on activity when the conformational equilibrium between open and closed state (K_C_) is maintained, i.e. the opening rate is scaled by the same factor. (b) Changing the rate constant for productive closing processes (ETM^O^ → ETM^C^ and ET^O^ →ET^C^) also has a small effect when K_C_ is preserved. The turnover is reduced when this step becomes rate limiting. In the orange curve the closing rate is reduced to 270 s^−1^, i.e. slower than substrate binding (Table S9) and the phosphotransfer step (500 s^−1^ in our model). (c) Changing the rate constant for unproductive closing without preserving the equilibrium does affect both the maximum turnover and substrate inhibition, with stronger inhibition for a faster closing rates. (d) In contrast, for the productive closing states accelerating domain closure has a positive effect on turnover. (e) Variation of the 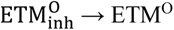 conversion rate. In the ATP and AMP-dependent simulations (Fig. 5b+c, Fig. S11a-d), a rate of 250 s^−1^ was presumed (blue). Increasing this rate by a factor of 10 (purple) relieves substrate inhibition largely, while a slower rate (0.1x, green) increases substrate inhibition.

**Table S1:** Kinetic parameters for the purified proteins in the high-throughput study.^a^

| **Mutation** | **$K_I$**<br>**($\mu\text{M}$)<sup>a</sup>** | **$K_M$**<br>**($\mu\text{M}$)<sup>a</sup>** | **ConSurf**<br>**grades<sup>b</sup>** | **Frequency of WT**<br>**amino acid<sup>c</sup> [%]** |
| --- | --- | --- | --- | --- |
| F19V | ($1.2 \cdot 10^5$) | 22.1 | 1 | 14 |
| I26D | 649 | 2.9 | 6 | 49 |
| I26V | 560 | 42.2 | 6 | 49 |
| P27S | 159 | 22.5 | 6 | 51 |
| P27V | 859 | 33.3 | 6 | 51 |
| I29E | 906 | 54.2 | 8 | 60 |
| I29M | ($1.7 \cdot 10^6$) | 46.9 | 8 | 60 |
| A37V | 412 | 62.6 | 4 | 46 |
| G42F | 1024 | 18.9 | 2 | 64 |
| L45S | 1437 | 19.8 | 6 | 63 |
| G46E | 1799 | 46 | 9 | 99 |
| G46S | 1577 | 35 | 9 | 99 |
| A66L | 398 | 27.5 | 1 | 15 |
| A66Y | 461 | 14 | 1 | 15 |
| K69E | 450 | 22.5 | 3 | 37 |
| E70A | 421 | 4.8 | 5 | 54 |
| E70S | 300 | 21 | 5 | 54 |
| A73E | 568 | 18.9 | 1 | 21 |
| N79V | 564 | 16.1 | 2 | 24 |
| L83F | 1472 | 29.9 | 8 | 75 |
| L83T | ($3.3 \cdot 10^6$) | 17.8 | 8 | 75 |
| L83Y | 581 | 25.6 | 8 | 75 |
| I90S | 528 | 159.6 | 3 | 29 |
| I90T | (2760) | 30.1 | 3 | 29 |
| I90V | 705 | 24.2 | 3 | 29 |
| I90Y | 1592 | 42.6 | 3 | 29 |
| P91E | ($3.4 \cdot 10^6$) | 141.9 | 2 | 31 |
| P91I | 419 | 14.2 | 2 | 31 |
| A93L | ($2.7 \cdot 10^6$) | 95 | 9 | 99 |
| A93T | (2010) | 23.6 | 9 | 99 |
| A93V | 1448 | 50.5 | 9 | 99 |
| A95V | 689 | 52.5 | 7 | 67 |
| M96I | 275 | 9.3 | 8 | 9 |
| E98K | 1755 | 24.1 | 2 | 38 |
| E98L | 1563 | 23.5 | 2 | 38 |
| E98N | 660 | 35.2 | 2 | 38 |
| E98V | 375 | 23.7 | 2 | 38 |
| A99E | 384 | 24.2 | 2 | 13 |
| G100M | 255 | 15.1 | 1 | 47 |
| G100T | 259 | 19.8 | 1 | 47 |
| G100V | 387 | 13 | 1 | 47 |
| I101A | 860 | 36.4 | 1 | 17 |
| I101V | 370 | 8.1 | 1 | 17 |
| N102D | 233 | 44.9 | 1 | 7 |
| V103M | 724 | 66.6 | 4 | 12 |
| Y105E | 305 | 3.9 | 1 | 11 |
| Y105T | 977 | 18.3 | 1 | 11 |
| Y105V | 1067 | 13.9 | 1 | 11 |
| V106F | (4.0·10<sup>6</sup>) | 75.7 | 8 | 81 |
| V106S | 625 | 21.9 | 8 | 81 |
| V117E | 1456 | 7 | 5 | 43 |
| A127K | 1545 | 138.3 | 3 | 11 |
| A127L | (2304) | 41 | 3 | 11 |
| A127S | 743 | 51.6 | 3 | 11 |
| P128I | 200 | 21.5 | 1 | 3 |
| P128T | 150 | 27.1 | 1 | 3 |
| K141D | 1209 | 15.4 | 4 | 45 |
| V142M | 484 | 22.1 | 1 | 28 |
| G144D | 584 | 49.4 | 5 | 70 |
| V148D | 602 | 85.5 | 1 | 27 |
| V148T | 728 | 30.2 | 1 | 27 |
| L153F | (3.3·10<sup>6</sup>) | 24.9 | 8 | 82 |
| L153T | 1729 | 16.4 | 8 | 82 |
| Q173M | 489 | 32.6 | 3 | 9 |
| A176Y | 223 | 13.4 | 2 | 25 |
| P177A | 632 | 81.1 | 7 | 82 |
| P177E | (2.1·10<sup>6</sup>) | 66.9 | 7 | 82 |
| L178M | (3.5·10<sup>6</sup>) | 115.1 | 7 | 72 |
| I179N | 661 | 12.1 | 5 | 38 |
| I179R | 1369 | 38.3 | 5 | 38 |
| G180S | 203 | 27.5 | 2 | 13 |
| K184S | 1276 | 48.9 | 1 | 23 |
| K184T | 290 | 24 | 1 | 23 |
| E187V | 560 | 36.4 | 1 | 8 |
| K192T | 545 | 88.7 | 2 | 43 |
| A207S | 1526 | 14.9 | 2 | 32 |
| A207T | 475 | 21.5 | 2 | 32 |
| A207V | 1107 | 26.3 | 2 | 32 |
| L209I | 330 | 7 | 6 | 24 |
| L209S | 331 | 8.3 | 6 | 24 |
| L209T | 1762 | 5.5 | 6 | 24 |
| I212A | 852 | 34.1 | 5 | 44 |
| I212L | 785 | 25.1 | 5 | 44 |
| I212T | 823 | 36.2 | 5 | 44 |
| I212V | 824 | 32.6 | 5 | 44 |
| L213S | 293 | 43.8 | 8 | 93 |

<sup>a</sup> $K_M$ and $K_I$ were derived by fits to Eq.1. Positions with mutants that result in a $K_I \leq 500 \mu\text{M}$ for AMP are marked in yellow, while those with $K_I \geq 1500$ are marked in green. Magenta refers to positions where both inhibited and uninhibited mutations are observed. Mutants with a very high value in $K_I$ ( $\geq 2000$ ) refer to non-inhibited proteins.
<sup>b</sup> ConSurf grades are conservation score derived based on a multiple sequence alignment (MSA) of 150 AK candidates using the ConSurf webserver. Higher numerical values correspond to higher conservation
<sup>c</sup> Frequency of the WT amino acid in the MSA

**Table S2:** Kinetic parameters for the activity of AK WT in dependence of ATP concentration (Fig. S3a).^a^

| ATP [M] | $v_{\max}$ [$s^{-1}$] | $K_M$ [$\mu M$] | $K_I$ [$\mu M$] |
| --- | --- | --- | --- |
| **1** | 487($\pm 41$) | 40.0($\pm 10.6$) | 2125($\pm 496$) |
| **2** | 499($\pm 34$) | 38.0($\pm 8.7$) | 2932($\pm 607$) |
| **3** | 544($\pm 35$) | 44.3($\pm 9.2$) | 3065($\pm 596$) |
| **4** | 526($\pm 22$) | 42.0($\pm 6.2$) | 4264($\pm 627$) |
| **5** | 586($\pm 32$) | 58.8($\pm 10.4$) | 4782($\pm 909$) |

<sup>a</sup> Fits are done according to a model of substrate inhibition involving dead-end complexes (Eq. 1)

**Table S3:**
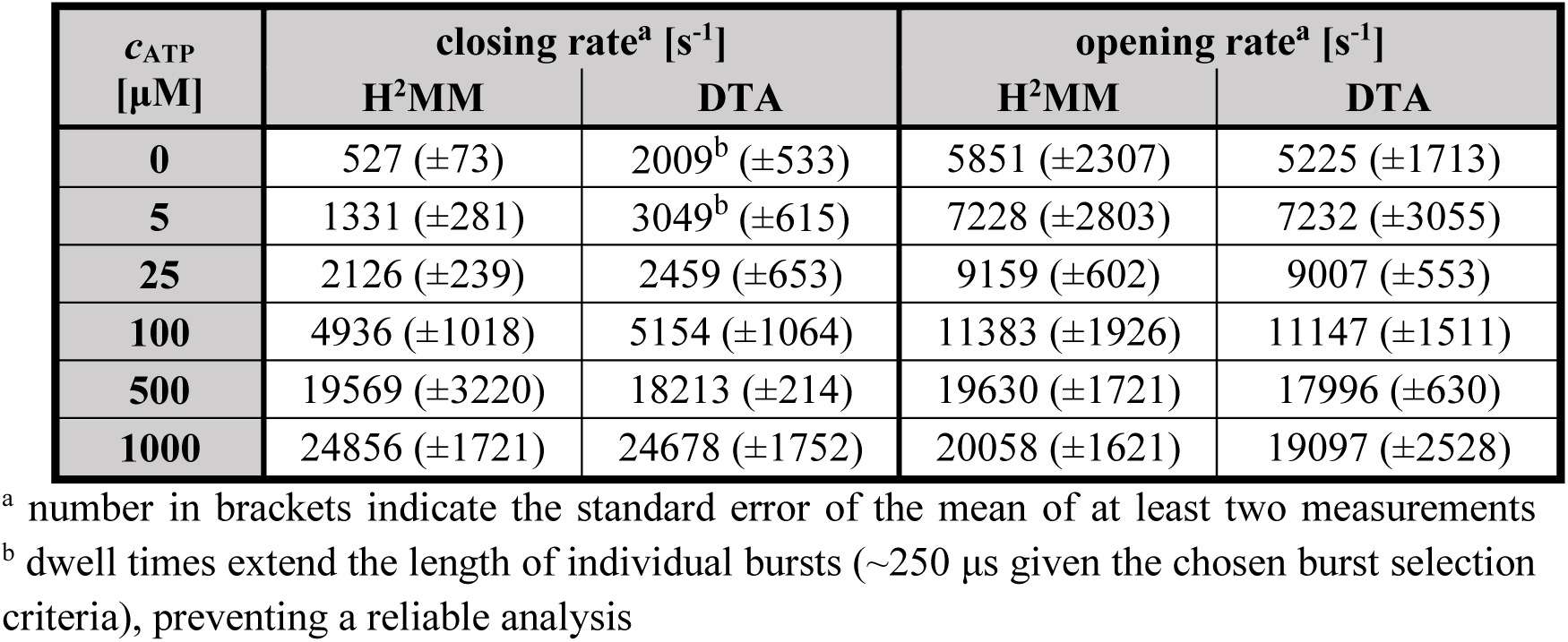
Comparison of transition rates obtained by H^2^MM vs. dwell-time analysis (DTA) for the WT protein.

**Table S4:** Parameters for fitted FCS curves of Figure S8 compared to the parameters obtained from the H^2^MM analysis.

|  | ATP<br>[μM] | 0 | 100 | 300 | 1000 |
| --- | --- | --- | --- | --- | --- |
| FCS<br>results <sup>a</sup> | N | 1.59±0.01 | 1.62±0.01 | 1.66±0.01 | 1.68±0.01 |
|  | p | 0.080±0.002 |  |  |  |
|  | T [μs] | 2.7±0.2 |  |  |  |
|  | K | 0.069±0.003 | 0.099±0.003 | 0.118±0.003 | 0.146±0.002 |
|  | τ<sub>c</sub> [μs] | 122±2 | 96±8 | 78±5 | 58±3 |
| τ<sub>c</sub> [μs] predicted based<br>on H<sup>2</sup>MM results <sup>b</sup> |  | 157±12 | 60±8 | 32±3 | 22±1 |

<sup>a</sup> Error bars represent the standard error of the fit
<sup>b</sup> Values are calculated based on the mean and the standard error of the mean of at least two measurements ( $\tau_c = 1/(k_{op} + k_{cl})$ )

**Table S5:** Substrate concentrations used in smFRET experiments.

| ***c*<sub>ATP</sub> fixed at 1 mM** |  |  | ***c*<sub>AMP</sub> fixed at 1 mM** |  |  |
| --- | --- | --- | --- | --- | --- |
| **AMP**<br>**[μM]** | **ADP**<br>**[μM]** | **Total conc.<sup>a</sup>**<br>**[μM]** | **ATP**<br>**[μM]** | **ADP**<br>**[μM]** | **Total conc.<sup>a</sup>**<br>**[μM]** |
| **1** | 6.2 | 1006 | **1\*10<sup>-6</sup>** | 2*10<sup>-3</sup> | 2*10<sup>-3</sup> |
| **2.5** | 9.9 | 1009 | **1\*10<sup>-5</sup>** | 0.02 | 0.02 |
| **5** | 13.9 | 1014 | **7.5\*10<sup>-5</sup>** | 0.05 | 0.05 |
| **10** | 19.7 | 1017 | **1\*10<sup>-3</sup>** | 0.2 | 0.2 |
| **25** | 31 | 1031 | **0.01** | 0.625 | 0.635 |
| **50** | 44 | 1044 | **0.025** | 1 | 1 |
| **100** | 62 | 1062 | **0.1** | 1.7 | 1.8 |
| **250** | 95 | 1095 | **0.5** | 3.7 | 4.2 |
| **400** | 123 | 1123 | **1** | 5.7 | 6.7 |
| **500** | 137 | 1137 | **2.5** | 8.4 | 10.9 |
| **750** | 150 | 1150 | **5** | 11.8 | 16.8 |
| **1000** | 160 | 1160 | **10** | 16.7 | 26.7 |
| **2000** | 230 | 1230 | **35** | 31 | 66 |
| **3000** | 327 | 1327 | **50** | 37.3 | 87.3 |
| **5000** | 417 | 1417 | **150** | 75 | 225 |
| **10000** | 576 | 1576 | **250** | 83 | 333 |
|  |  |  | **1000** | 160 | 1160 |
|  |  |  | **2000** | 230 | 2230 |
|  |  |  | **5000** | 417 | 5417 |
|  |  |  | **10000** | 576 | 10576 |

<sup>a</sup> total concentration of substrates that can bind to the LID domain, i.e. ATP and ADP

**Table S6:** Fitted parameters for the ATP-dependent closure (Figure 3).^a^

|  | WT |  | L107I |  | L82V |  | F86W |  |
| --- | --- | --- | --- | --- | --- | --- | --- | --- |
| AMP | + | - | + | - | + | - | + | - |
| $C_{50,ATP}$ [μM] | 176<br>(±17) | 9.0<br>(±0.9) | 127<br>(±32) | 6.9<br>(±0.8) | 20<br>(±2.3) | 0.55<br>(±0.04) | 223<br>(±24) | 172<br>(±49) |
| $Occ_{EX}$ [%] | 12.8<br>(±1.0) | 15.6<br>(±1.0) | 13.0<br>(±1.7) | 15.3<br>(±1.0) | 15.1<br>(±1.2) | 21.6<br>(±0.7) | 9.9<br>(±1.0) | 11.5<br>(±2.0) |
| $Occ_{EXT}$ [%] | 62.5<br>(±1.1) | 57.1<br>(±0.6) | 56.9<br>(±1.8) | 64.2<br>(±0.9) | 78.0<br>(±1.1) | 78.4<br>(±0.7) | 65.3<br>(±1.4) | 55.8<br>(±2.3) |

<sup>a</sup> Columns/mutants are color-coded according to color-palette used in the main text**Table S7:** Parameters for the impact of AMP on LID domain dynamics (Figure 4a-c) according to the model described in Supplementary Note 1.

**Table S7:**
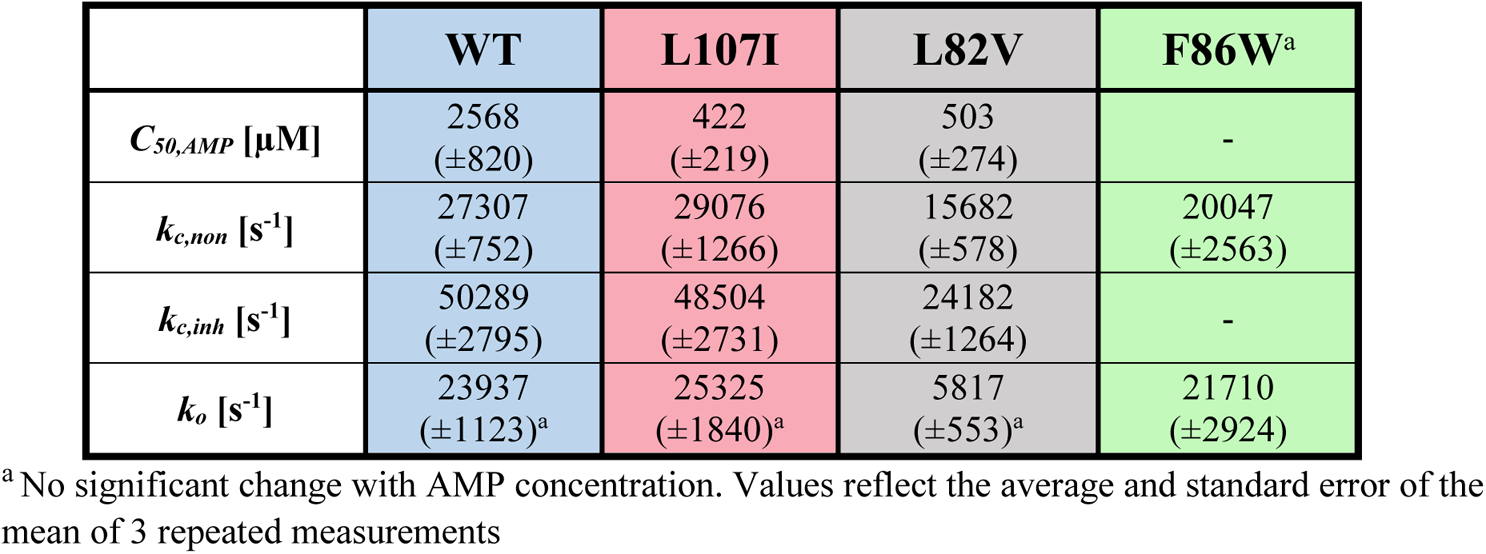
Parameters for the impact of AMP on LID domain dynamics (Figure 4a-c) according to the model described in Supplementary Note 1.

|  | WT | L107I | L82V | F86W<sup>a</sup> |
| --- | --- | --- | --- | --- |
| $C_{50,AMP}$ [μM] | 2568<br>(±820) | 422<br>(±219) | 503<br>(±274) | - |
| $k_{c,non}$ [s<sup>-1</sup>] | 27307<br>(±752) | 29076<br>(±1266) | 15682<br>(±578) | 20047<br>(±2563) |
| $k_{c,inh}$ [s<sup>-1</sup>] | 50289<br>(±2795) | 48504<br>(±2731) | 24182<br>(±1264) | - |
| $k_o$ [s<sup>-1</sup>] | 23937<br>(±1123)<sup>a</sup> | 25325<br>(±1840)<sup>a</sup> | 5817<br>(±553)<sup>a</sup> | 21710<br>(±2924) |

<sup>a</sup> No significant change with AMP concentration. Values reflect the average and standard error of the mean of 3 repeated measurements

**Table S8:** Opening and closing rates used in the simulation of experimental activity patterns.

|  |  | WT |  | L107I |  | L82V |  | F86W |  |
| --- | --- | --- | --- | --- | --- | --- | --- | --- | --- |
| rate constants<br>[$\times 10^3 \text{ s}^{-1}$] | | O→C | C→O | O→C | C→O | O→C | C→O | O→C | C→O |
|  | E | 0.5 | 6 | 2.2 | 17 | 1.5 | 10 | 0.34 | 3.9 |
|  | EM | 2.6 | 13 | 7.5 | 38 | 16 | 37 | 0.36 | 3.6 |
|  | ET<sup>a</sup> | 27 | 24 | 30 | 25 | 16 | 6 | 20 | 22 |
|  | ETM<sub>inh</sub> | 50 | 24 | 49 | 25 | 24 | 6 | 20 | 22 |
|  | ETM<sup>a</sup> | 27 | 24 | 30 | 25 | 16 | 6 | 20 | 22 |

<sup>a</sup>The single molecule experiments cannot distinguish between ET and ETM.

**Table S9:** Nucleotide binding rates used in the simulation of experimental activity patterns.

|  | WT |  | L107I |  | L82V |  | F86W |  |
| --- | --- | --- | --- | --- | --- | --- | --- | --- |
| productive binding rates [$\mu\text{M}^{-1} \text{ s}^{-1}$] | ATP | AMP | ATP | AMP | ATP | AMP | ATP | AMP |
|  | 9.31 | 5.24<sup>a</sup> | 12.49 | 6.29<sup>a</sup> | 3.62 | 13.13<sup>a</sup> | 0.68 | 0.07 |

<sup>a</sup>In the case of substrate inhibition (WT, L107I and L82V with AMP), parameters were approximated using Eq. (1).

**Table S10:**
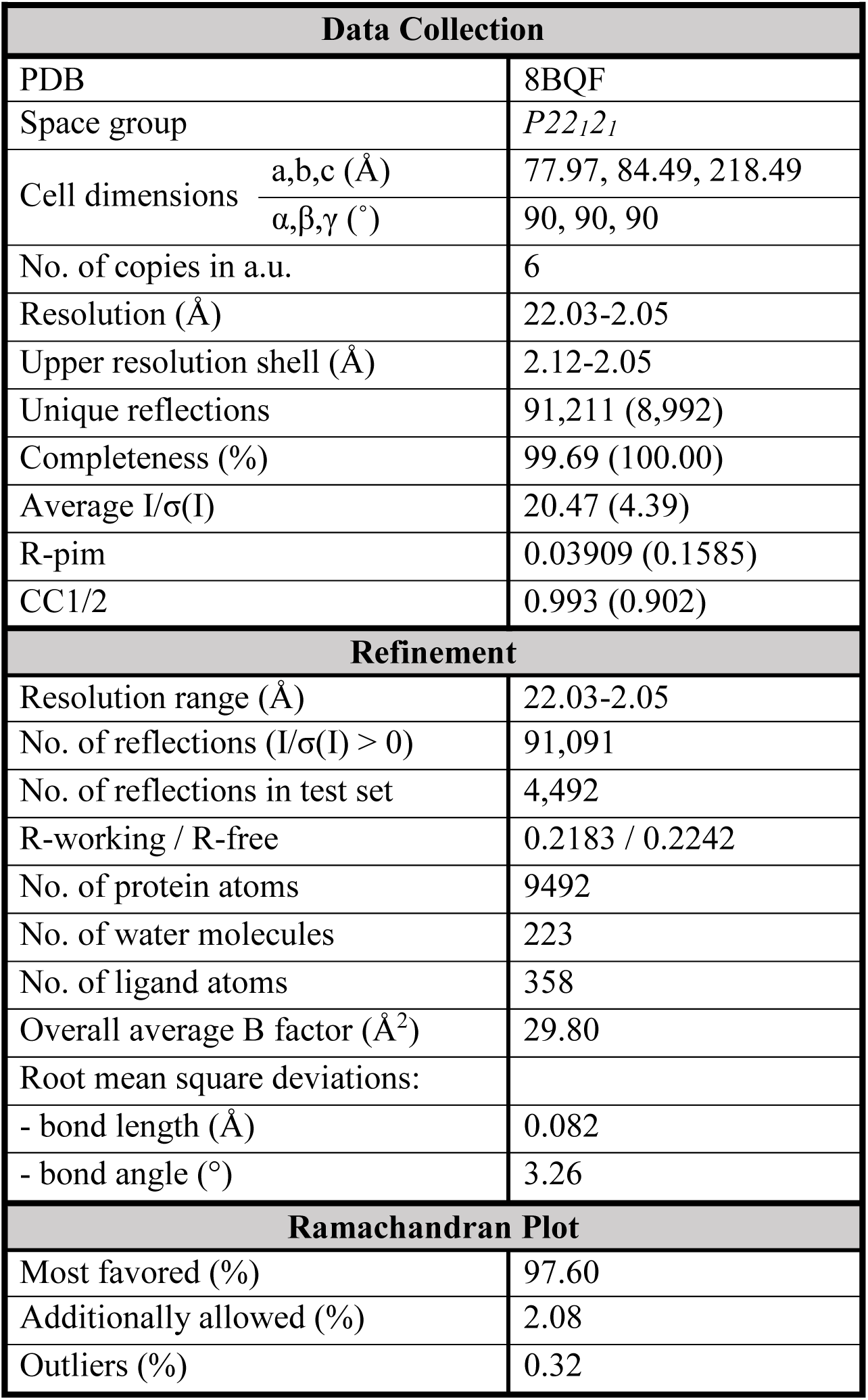
Data collection and refinement statistics for the crystal structure of AK L107I in complex with Ap_5_A.

